# A unifying model for discordant and concordant results in human neuroimaging studies of facial viewpoint selectivity

**DOI:** 10.1101/2023.02.08.527219

**Authors:** Cambria Revsine, Javier Gonzalez-Castillo, Elisha P Merriam, Peter A Bandettini, Fernando M Ramírez

## Abstract

Our ability to recognize faces regardless of viewpoint is a key property of the primate visual system. Traditional theories hold that facial viewpoint is represented by view-selective mechanisms at early visual processing stages and that representations become increasingly tolerant to viewpoint changes in higher-level visual areas. Newer theories, based on single-neuron monkey electrophysiological recordings, suggest an additional intermediate processing stage invariant to mirror-symmetric face views. Consistent with traditional theories, human studies combining neuroimaging and multivariate pattern analysis (MVPA) methods have provided evidence of view-selectivity in early visual cortex. However, contradictory results have been reported in higher-level visual areas concerning the existence in humans of mirror-symmetrically tuned representations. We believe these results reflect low-level stimulus confounds and data analysis choices. To probe for low-level confounds, we analyzed images from two popular face databases. Analyses of mean image luminance and contrast revealed biases across face views described by even polynomials—i.e., mirror-symmetric. To explain major trends across human neuroimaging studies of viewpoint selectivity, we constructed a network model that incorporates three biological constraints: cortical magnification, convergent feedforward projections, and interhemispheric connections. Given the identified low-level biases, we show that a gradual increase of interhemispheric connections across network layers is sufficient to replicate findings of mirror-symmetry in high-level processing stages, as well as view-tuning in early processing stages. Data analysis decisions—pattern dissimilarity measure and data recentering—accounted for the variable observation of mirror-symmetry in late processing stages. The model provides a unifying explanation of MVPA studies of viewpoint selectivity. We also show how common analysis choices can lead to erroneous conclusions.

## 1. INTRODUCTION

The ability of humans to recognize faces regardless of viewpoint remains a topic of intensive research. From a computational perspective (Marr, 1982), viewpoint-invariant recognition is challenging due to vast variation among images corresponding to the same identity. Theoretical work suggests that exploiting object symmetries, such as bilateral symmetry of the head, might substantially reduce the complexity of this task (Vetter et al., 1994; Poggio and Anselmi, 2016; Leibo et al., 2017). Current research aims to understand how and where object symmetries are represented in the brain to aid viewpoint-invariant recognition.

In macaques, face recognition is thought to be supported by a hierarchically-organized system of face-selective patches along the ventral visual pathway (Moeller et al., 2008; Tsao et al., 2008). Freiwald and Tsao (2010) characterized the tuning properties of neurons recorded from three stages of this hierarchy. Neurons in the middle lateral/fundus face patch (ML/MF) maximally responded to faces in one preferred view. In contrast, anterior lateral (AL) neurons exhibited bimodal tuning functions maximally responsive to mirror-symmetric face views. Finally, the anteromedial face patch (AM) exhibited virtual viewpoint-invariance. It was proposed that AL may be an intermediate mirror-symmetrically tuned processing stage along the face-processing hierarchy.

Motivated by the goal of finding evidence of viewpoint-compartmentalization, as observed in macaques, several studies have attempted to characterize the form of view-tuning of neural populations in human face-selective areas with a combination of functional magnetic resonance imaging (fMRI) and multivoxel pattern analyses (MVPA). These studies measured blood-oxygen-level-dependent (BOLD) fMRI responses in the occipital face area (OFA), fusiform face area (FFA), posterior superior temporal sulcus (pSTS), and early visual cortex (EVC), and characterized these areas as predominantly view-tuned or mirror-symmetric (Figure 1). While all studies detailed in Table 1 observed evidence of view-tuning in EVC, conclusions in increasingly anterior face-selective areas proved increasingly harder to reconcile. For example, while some studies concluded that neurons in human FFA are tuned to a single preferred view—like the macaque ML/MF face patches, other studies concluded that FFA is mirror-symmetrically tuned—like the AL face-patch.

**Figure 1.**
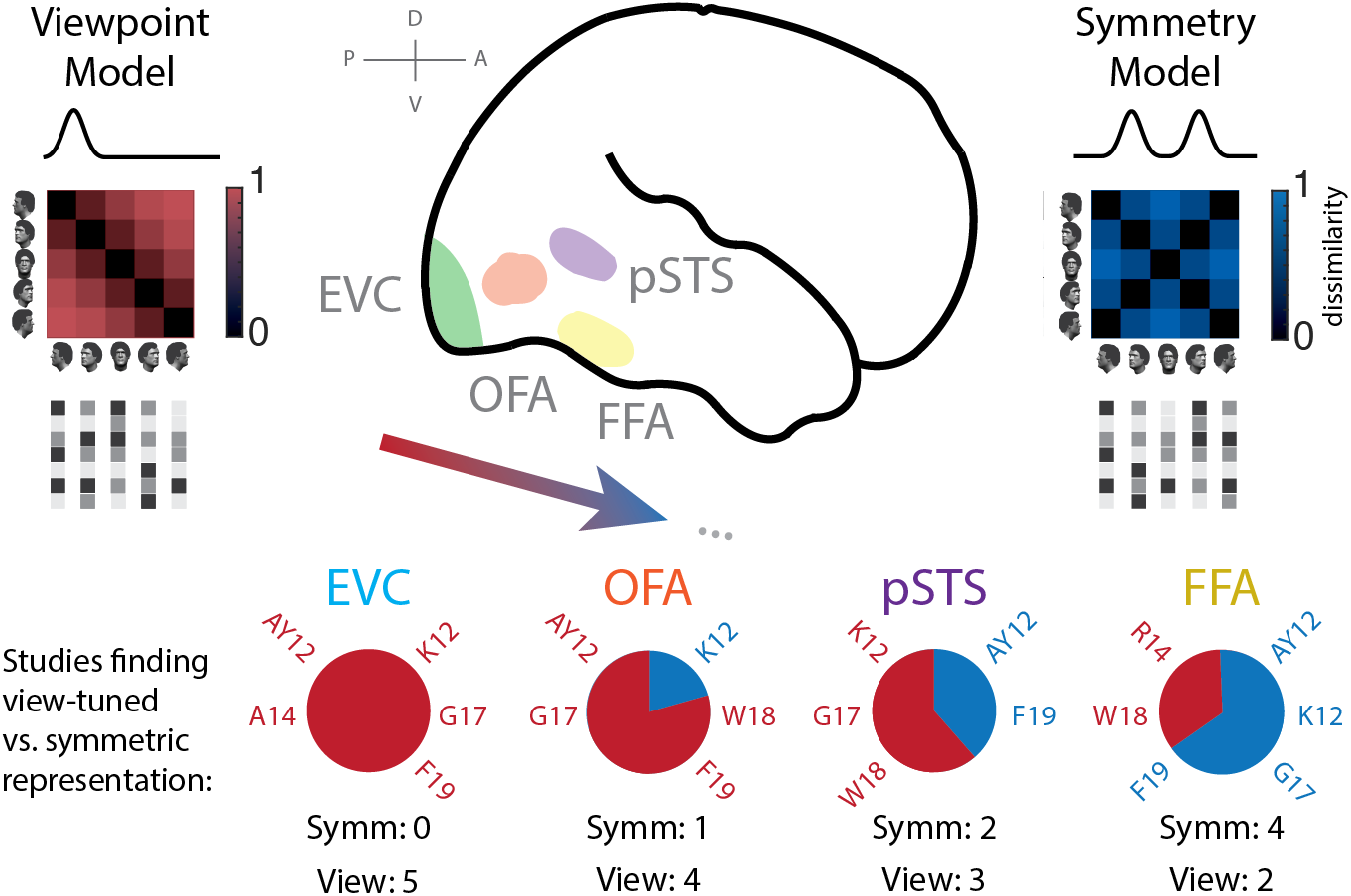
Commonalities and inconsistencies across fMRI-MVPA studies investigating viewpoint representations in humans. *a)* Four regions of interest are depicted on a sagittal view of the brain. In early visual cortex (EVC, shown in green), 5/5 studies reported marked view-tuning, as depicted by the dissimilarity matrix shown in red (Viewpoint model) and the unimodal neural tuning function shown immediately above. In OFA (in orange), 5/5 studies reported evidence of view-tuning. One study, however, reported additional evidence of some degree of mirror-symmetry, represented by the blue dissimilarity matrix (Symmetry model) and bimodal tuning function immediately above. In pSTS (in purple), 5/5 studies reported evidence of view-tuning, with 2/5 observing some degree of mirror-symmetry. Finally, in the FFA (in yellow), 6/6 studies reported evidence of view-tuning, while 4/6 of these studies also reported evidence of mirror-symmetry, ranging from weak to strong. In sum, while marked view-tuning was consistently observed in posterior brain regions, mirror-symmetry was inconsistently observed, albeit with increasing frequency, in increasingly anterior areas along the ventral stream. OFA: occipital face area, pSTS: posterior superior temporal sulcus, FFA: fusiform face area.

**Table 1.**

| Study Name | Abbreviation | Task Design | Pattern Estimation Method | MVPA Type | RSA Distance Measure | Data Re-Centering | ROI Def. (Bilateral/Unilateral) |
| --- | --- | --- | --- | --- | --- | --- | --- |
| Axelrod & Yovel, 2012<br>*Exp. 1* | AY12a | *No fixation*<br>One-back, identity | *none*<br>Concatenation of raw† intensity values for each condition within runs | Multiclass linear SVC, spatiotemporal patterns | Error rates | *Yes*<br>a. Time series z-scored<br>b. Patterns demeaned across voxels | Bilateral<br>EVC, FFA; right OFA, pSTS |
| Axelrod & Yovel, 2012<br>*Exp. 2* | AY12b | *Yes fixation*<br>color change detection of fixation spot | *none*<br>Concatenation of raw† intensity values for each condition within runs | Multiclass linear SVC, spatiotemporal patterns | Error rates | *Yes*<br>a. Time series z-scored<br>b. Patterns demeaned across voxels | Bilateral<br>EVC; right OFA, FFA |
| Kietzmann et al., 2012 | K12 | *Yes fixation*<br>One-back, identity | Averaging for each block of percent signal changes of TRs 5-12 (10-24 sec.) | RSA | Correlation | *Yes*<br>a. Time series z-scored<br>b. Regressed out mean spatial pattern | Bilateral<br>EVC, OFA, FFA |
| Anzellotti et al., 2014 | A14 | *No fixation*<br>Identity detection | GLM with canonical HRF | 1. RSA<br>2. Linear SVC | a. Correlation | *No* | Right<br>EVC, OFA, FFA |
| Ramírez et al., 2014 | R14 | *Yes fixation*<br>luminance change detection | GLM with canonical HRF | 1. RSA<br>2. Linear SVC | a. Correlation<br>b. Euclidean | *No* | Bilateral<br>EVC; right OFA, pSTS, FFA |
| Guntupalli et al., 2017 | G17 | *No fixation*<br>One-back, identity | GLM with canonical HRF | 1. RSA<br>2. Linear SVC, spatiotemporal patterns | a. Correlation<br>b. Classification accuracy | *No* | Bilateral<br>EVC, OFA, FFA; right pSTS |

SVC: Support Vector Classification; GLM: General Linear Model; HRF: Hemodynamic Response Function; EVC: Early Visual Cortex; FFA: Fusiform Face Area; OFA: Occipital Face Area; pSTS: posterior Superior Temporal Sulcus. (†) fMRI data were motion corrected, spatially normalized, smoothed, and detrended.

How might such inconsistencies emerge between fMRI studies using similar stimuli and experimental designs? One possibility lies in different analysis methods, including the choice whether to demean the data, and what dissimilarity measure to use in Representational Similarity Analyses (RSA) (Kriegeskorte et al., 2008). We previously observed that studies reporting mirror-symmetry either demeaned the data prior to RSA analyses using the correlation distance to measure pattern dissimilarities, or relied on measures related to the Euclidean distance. We also noted that stronger overall responses for frontal than profile face-views in visual cortex have been reported in multiple fMRI studies (reviewed in Ramírez, 2018). Critically, regardless of the form of viewpoint selectivity of the underlying neuronal populations, stronger responses for frontal than lateral face views will lead to mirror-symmetry with analysis pipelines that either demean the data prior to RSA or use the Euclidean distance to measure pattern dissimilarities (Ramírez, 2017). We recently also showed that stimuli from previous studies exhibit low-level feature imbalances across face-views consistent with the reported overrepresentation (Ramírez et al., 2020). Building on these observations, geometrical reasoning led us to hypothesize that interhemispheric crossings in the brain might explain common and inconsistent trends observed across neuroimaging studies of viewpoint selectivity (see Figure 2). We reasoned that the influence of low-level stimulus features (e.g., image luminance and contrast) in a visual area must depend on the relative strength of contra- and ipsilateral hemifield responses, and therefore qualitatively different biases in signal-strength across conditions along the visual hierarchy would differently interact with data-analysis choices.

**Figure 2.**
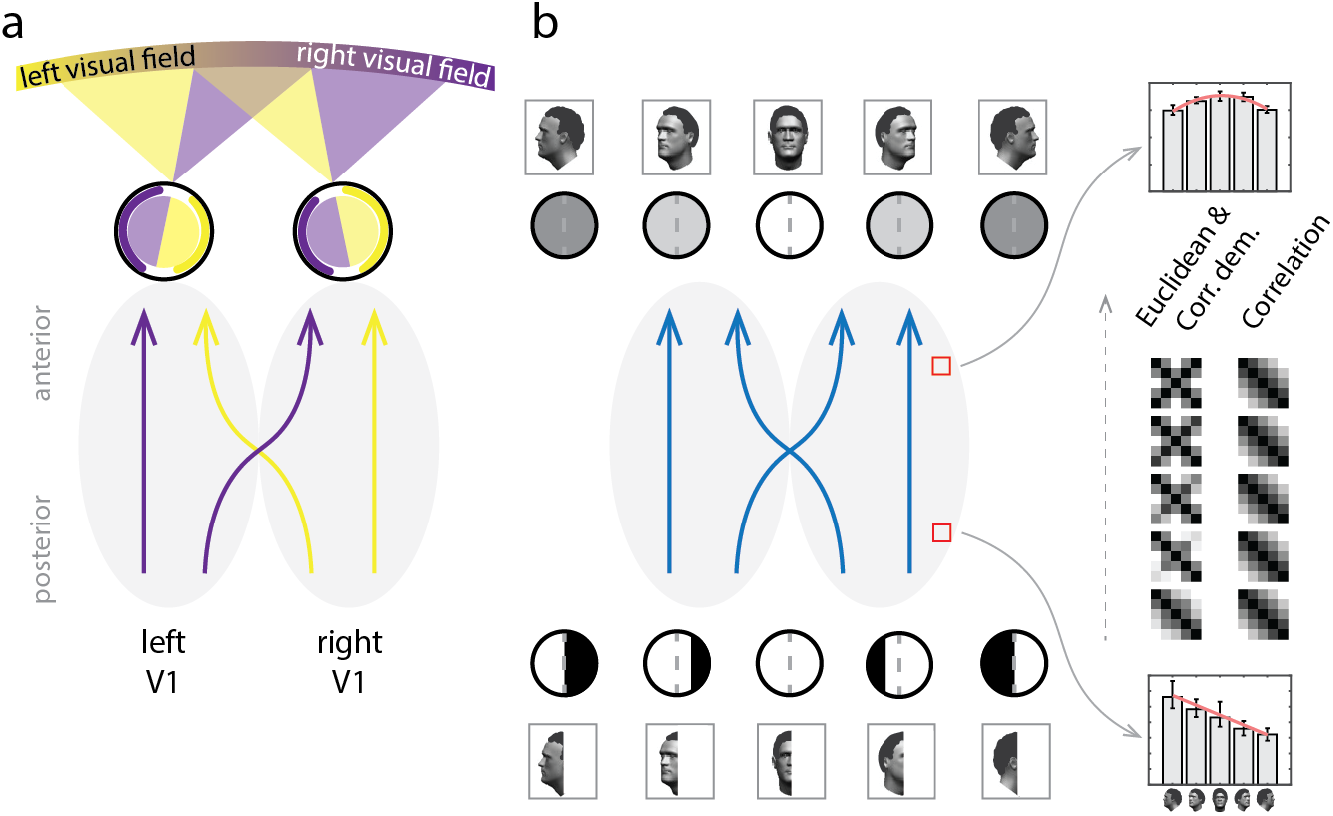
Proposed account of commonalities and inconsistencies across fMRI-MVPA studies. *a)* Schematic of visual hemifields and their mapping onto cerebral hemispheres, axial view. Locations in left and right visual fields map onto primary visual cortex of the contralateral cerebral hemisphere. A known property of visual cortex is increasingly bilateral hemifield representations, due to interhemispheric connections, as one proceeds along the visual hierarchy. This property is central to the model proposed here. *b)* (Low-level image properties) + (interhemispheric crossings) ≈ RSA results. Faces in different views exhibit different distributions of luminance and contrast. These properties have been found to exhibit symmetric distributions about the frontal view. Full circles (top row) indicate the mean luminance of the image of the face view shown immediately above. Anterior brain areas, which integrate input from both hemifields, are expected to exhibit quadratic (i.e., symmetric) biases of the form illustrated in the bar-plot shown at the top-right. In contrast, responses across views for half-images (bottom row) are expected to exhibit antisymmetric biases. Black and white circles indicate the rough luminance distribution of each half-image for each face view—note the dark hair and bright skin. For right V1, this would imply a roughly linear (i.e., antisymmetric) trend if responses were proportional to the luminance of the left side of the stimulus (see bar-plot at bottom right). If RSA outcomes reflect such trends for luminance and contrast, as we propose here, then dissimilarity matrices at earlier processing stages would exhibit marked view-tuning regardless of pattern dissimilarity measure. In turn, representations in later processing stages would exhibit mirror-symmetry either with the Euclidean distance, or angular distances (e.g., correlation distance) *only* if the data are mean-centered across conditions prior to RSA. Instead, for RSA with the correlation distance, a viewpoint-specific representation would be expected throughout cortex, as shown by the dissimilarity matrices in the rightmost column.

To test this hypothesis, we constructed a network model incorporating three biologically-motivated constraints: convergent feedforward projections, interhemispheric crossings increasing in frequency along the processing hierarchy, and foveal cortical magnification. Connections between network units were otherwise randomly generated, and hence not explicitly tuned to viewpoint information. We evaluated this minimally-structured model in its capacity to reproduce observations across fMRI-MVPA studies when given as input images from two popular face databases. The model explained why conclusions in EVC are consistent across studies regardless of data recentering and RSA distance measure, while results in higher-tier areas are sensitive to these analysis choices.

## 2. METHODS

All simulations and statistical analyses were implemented using MATLAB (R2018b) and executed in Biowulf, the high-performance computing facility at NIH, as well as tested on a MacBook Pro (2016, 16 GB RAM) running MacOS (10.15.7). The implemented network architecture consisted of eight layers and considered exclusively feedforward connections between units in adjacent layers. Two further constraints on the architecture were (i) cortical magnification (CM) of central image locations in the input layer, and (ii) a gradual increase in the number of interhemispheric projections in subsequent levels of the hierarchy. A preliminary version of this study was published in abstract form (Revsine et al., 2020).

In the simulations described here, when a specific instantiation of the model receives an image as input, it outputs a set of distributed response patterns—one per network layer. We obtained patterns associated with faces belonging to various identities, each presented in five viewpoints (see Section 2.1 for details). Response patterns in each network-layer were then subjected to three variants of Representational Similarity Analysis (RSA). A first factor distinguishing these variants was the distance measure used to describe the dissimilarity between activity patterns. We computed dissimilarity matrices according to the Euclidean (RSA_Euc_) and the correlation distance (RSA_corr_). A second distinguishing factor was the choice whether to demean the simulated response patterns prior to computing pattern dissimilarities. By “demeaning” throughout this paper we refer to the practice of subtracting in each measurement channel—e.g., fMRI voxel—the mean activity-level observed across conditions from that associated with each condition. Only RSA using the correlation distance was conducted both on native and demeaned activity patterns (see Section 2.4.2 for details). We chose these three analysis variants because they reflect key differences in the analysis pipelines used by previous studies. To explore the impact of network connectivity density on pattern analyses, our model included a parameter that controlled the number of connections received by each network unit.

Below, we describe in further detail the (i) images provided as input to the model, (ii) network architecture, (iii) two model variants explored, and (iv) three pattern analysis variants conducted on the distributed response patterns provided as output by the model.

### 2.1 Input images

Images provided as input to the model were obtained from two popular databases: Karolinska Directed Emotional Faces (KDEF) (Lundqvist et al., 1998) and Radboud Faces Database (RaFD) (Langner et al., 2010). These databases have been used by previous studies investigating viewpoint-invariant face recognition, and RaFD specifically used in studies combining fMRI and MVPA (Weibert et al., 2018; Flack et al., 2019). Both databases include multiple individuals photographed in the same five viewpoints: −90° (left-profile view), −45°, 0°, 45°, and 90° (right-profile view). The RaFD database contains images of 57, and the KDEF database of 70, adult individuals. Photographs of two individuals from the KDEF database were excluded because they exhibited lighting conditions inconsistent with the remaining identities (mean luminance > 4 SD away from the mean). Both databases include images of each identity displaying various emotions and eye-gaze directions. Only faces with neutral expressions and consistent eye-gaze and head-direction were provided as input to the model.

Images were converted from RGB to grayscale values (range: 0 to 255) and resized to ensure the height of the head measured on average 8.7° of visual angle, well within the range used in previous studies [range: 4.2° to 12.5° (vertical dimension)]. Image size was specified with respect to the input array of the network model (441 pixels × 441 pixels), which spanned 12.1° × 12.1° of visual angle. The center of the fovea was located at the central pixel of the array. Images were centered and overlaid on a uniform gray background. An example face image is shown in Figure 4. Note that face images throughout this manuscript are used for illustration only, computer generated, and not photographs from the analyzed databases.

### 2.2 Network architecture

The network consists of eight hierarchically organized layers, and is comprised of two hemispheres. Each network layer consists of multiple units; 4096 units in layer 1 (2048 per hemisphere), and 1024 units in each of the remaining layers (2 to 8) (512 per hemisphere). Units project exclusively onto units in the subsequent layer. Each network layer is intended to correspond to a different processing stage along the ventral visual processing stream. While early visual areas can be roughly mapped onto the first two or three layers of the network, high-level face-selective areas such as OFA and FFA are assumed to correspond to later layers of the network, with OFA and pSTS presumably in a hierarchical level preceding that of FFA (but see Rossion et al., 2011). However, we do not suggest an explicit mapping between brain areas and layers of our network model.

One prominent aspect of our network architecture is that it considers distinct left and right network hemispheres, emulating the structure of the brain (Figures 2 and 3). Connections between units in consecutive layers originate from either the ipsilateral or contralateral hemisphere of the previous layer. In successive layers of the network, connections are increasingly likely to cross hemispheres, reflecting the increasing probability of neural projections crossing through the corpus collosum and anterior commissure at successive stages of the visual hierarchy (Berlucchi, 2014). This aspect of the model is supported by a well-documented increase of receptive field (RF) sizes along the ventral visual hierarchy, such that RFs in higher-tier areas tend to be large, usually include the fovea, and cross the vertical meridian into the ipsilateral hemifield (Gross et al., 1972; Desimone et al., 1984; Rolls, 2012). Human fMRI measurements are consistent with this concept (Tootell et al., 1998; Hemond et al., 2007; Dumoulin and Wandell, 2008; Henriksson et al., 2008; Amano et al., 2009). The RF of each unit in layer 1 is defined as the collection of image locations that provide input to that unit. Units in the left hemisphere of layer 1 receive input almost entirely from pixel locations in the right hemifield of the image, and vice versa for units in the right hemisphere. A second biological property incorporated by the model is cortical magnification (CM). Image locations towards the center of each image are sampled with markedly increased probabilities, following the cortical magnification function described in Duncan and Boynton (2003).

**Figure 3.**
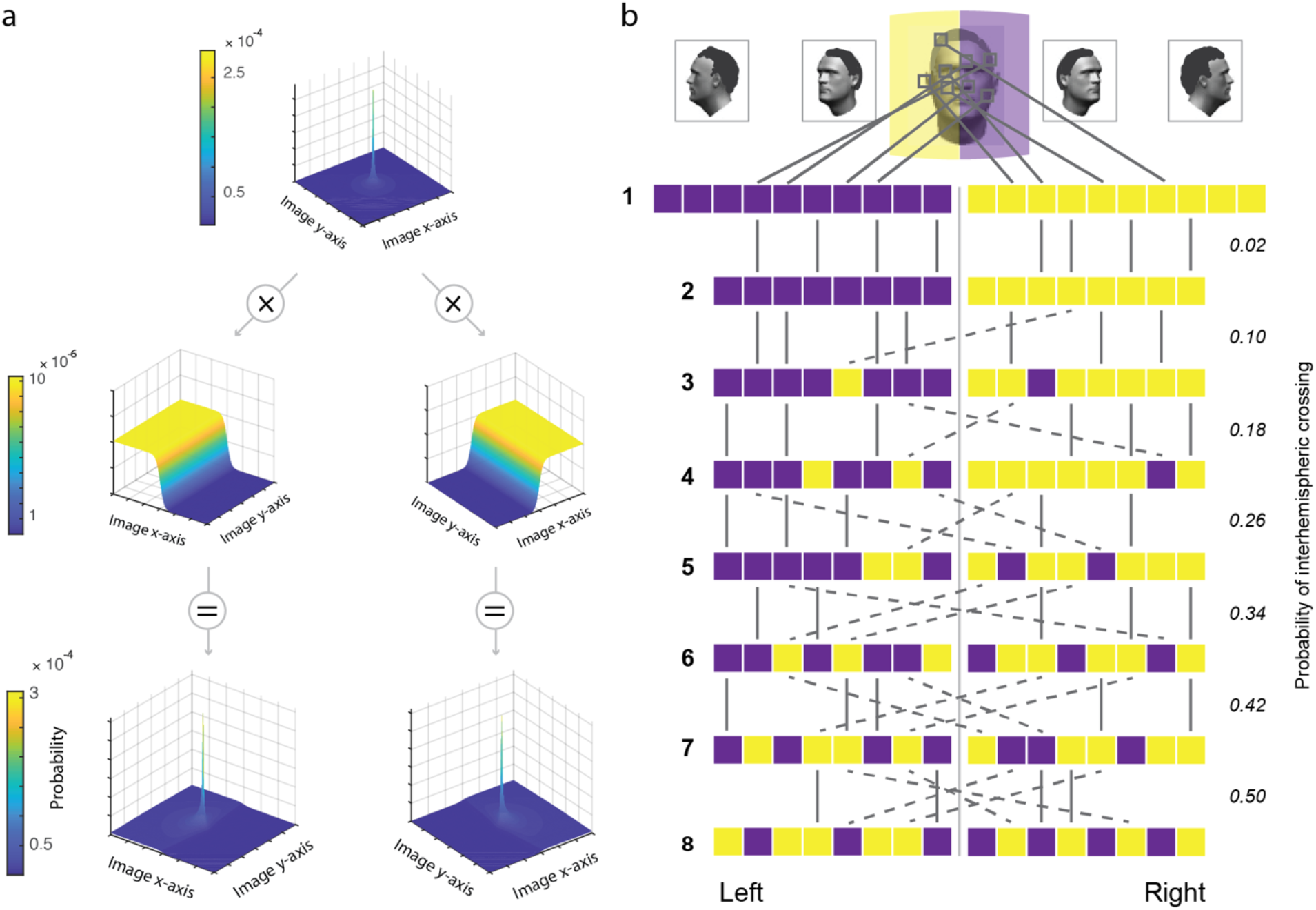
Feedforward, randomly-connected, two-hemisphere network architecture. *a*) Probability distributions used to specify image locations sampled by Layer 1 units. Top row: Distribution used to model cortical magnification (CM) of central image locations in V1. Middle row: Distributions used to model left-hemifield (LH) and right-hemifield (RH) representations. Bottom row: Product of CM with LH and RH distributions. These distributions served to specify, by random sampling, image locations providing input to units in each hemisphere of Layer 1. *b*) Full network, consisting of eight layers; Layer 1 (4096 units) and Layers 2-8 (each 1024 units). Feedforward connections between units in consecutive layers define this architecture. Input to the left hemisphere of Layer 1 (shown in purple) originates from receptive fields located almost exclusively on the right side of each image. The opposite is observed for the right hemisphere of Layer 1 (in yellow). Ipsilateral projections within each network hemisphere are indicated by solid lines. Contralateral projections are indicated by dashed lines. The probability of a contralateral projection increases in steps of 0.08, beginning at 0.02 between layers 1 and 2, and reaching 0.5 between layers 7 and 8.

#### 2.2.1 Probability distributions: layer 1

To specify the input received by each layer 1 network unit, we randomly sampled pixel locations from the 441 x 441 array assumed in our model to represent input to the retinae. The exact number of pixel-locations providing input to a particular unit in layer 1 was specified by a parameter controlling the connectivity density of the network (see Section 2.2.3 for details). For clarity, we will first explain the random process by which we specify the pixel locations providing input to each layer 1 unit, which comes almost exclusively from the contralateral hemifield. To this aim, we relied on two probability distributions—termed cortical magnification left hemifield (CM-LH) and cortical magnification right hemifield (CM-RH). Each of these distributions was tailored to reflect two properties of human primary visual cortex (V1), namely, (i) cortical magnification and (ii) separate hemifield representations. CM-LH and CM-RH were obtained by combining, in each case, a distribution used to specify the preferred hemifield of a unit, with a second distribution used to model the desired overrepresentation of the fovea (see Figure 3*a*). CM-LH and CM-RH are mirror-symmetric versions of each other. Importantly, both distributions exhibit markedly increased probabilities at locations of the input array corresponding to the fovea. These probability distributions were used to specify the multinomial probability assignments according to which pixel-locations were then randomly drawn to define the RF of each unit.

In more detail, as a first step, a two-dimensional distribution was specified using the following formula for the linear cortical magnification factor of V1 described in Duncan and Boynton (2003):

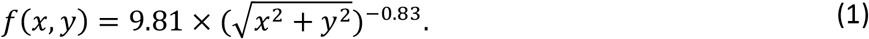

In this formula, 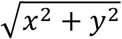 denotes eccentricity from the origin, which is in our model at the center of the fovea. Because the value of this function at the origin tends to infinity, we arbitrarily set this value to 100. The distribution was then normalized to ensure the sum of discrete probabilities was equal to one. This distribution models heightened spatial resolution at the foveal confluence. Probabilities exponentially decrease as a function of distance from the fovea. We will refer to this distribution as the cortical magnification (CM) distribution (Figure 3*a*, top row).

To model the existence of a left and right visual hemifield representation in V1, we then defined two 2D distributions, one corresponding to the left hemifield, the other to the right hemifield (Figure 3*a*, middle row). To define these distributions, we relied, respectively, on the following logistic functions:

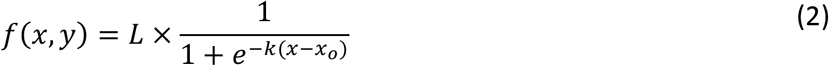

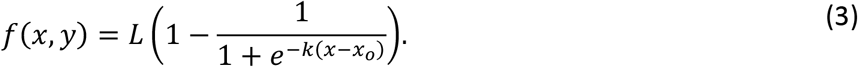

In these functions, *L* specifies the maximum value of the curve, *x_o_* the value of the midpoint along the x-axis, and *k* the logistic growth rate. Here, *L* was set to 10, *x_o_* to 0, and *k* to 4. These values achieve a steep transition across the vertical meridian from low to high probabilities. Probability densities were maximally concentrated on the left hemifield for the left-hemifield distribution, and on the right hemifield for the right-hemifield distribution. The output values of functions (2) and (3), when evaluated on any arbitrary pair of x- and y-values, are independent of the value of y, and hence the resulting probability distributions can be understood as a concatenation of 1D logistic distributions along the y-dimension (see Figure 2*a*, middle row).

Finally, the CM distribution and the left-hemifield logistic distributions were pointwise multiplied and normalized to ensure discrete probabilities summed to one. We term the resulting distribution CM-LH (Figure 3*a*; bottom row, left). This distribution is characterized by probabilities maximally concentrated at the center of the input array, a property inherited from the CM distribution, and heightened probabilities in the left hemifield, inherited from the left-hemifield logistic distribution. In a separate but otherwise identical procedure, CM was pointwise multiplied by the right-hemifield logistic distribution and normalized to produce the CM-RH distribution (Figure 3*a*; bottom row, right).

To explore the relevance of incorporating cortical magnification into the model, we tested an additional variant excluding this specific component from the model. In this reduced model, only the left-hemisphere and right-hemisphere logistic distributions were used to specify input projections to units in layer 1.

#### 2.2.2 Network architecture: layers 2 – 8

Units in layers 2-8 of the network, like layer 1 units, receive feedforward input exclusively from the immediately preceding processing stage. Unlike layer 1 units, however, which receive projections almost exclusively from the contralateral hemifield of the input image, inputs to units in the remaining layers are specified according to a binomial distribution. The latter distribution is used to determine whether each projection received by units beyond layer 1 originate from the ipsilateral or contralateral hemisphere in the preceding network layer. As was the case in layer 1, the number of input connections received by each unit is again controlled by the density parameter *d* (see Section 2.2.3). By parametrically changing the binomial probability *p* over successive layers, we enforce on the model a gradual increase in the frequency of interhemispheric connections along the network hierarchy.

When defining connections between layer 1 and 2, binomial probabilities were set to a value of 0.02. This implies that there is a 0.02 probability that the feedforward projection to that layer 2 unit corresponds to a contralateral projection, and a 0.98 probability that it corresponds to an ipsilateral connection. Binomial probabilities of a contralateral connection in subsequent layer transitions, as specified by parameter *p*, were sequentially updated in steps of 0.08 until reaching a probability of 0.5 at the transition between layers 7 and 8 (see Figure 3*b*). The exact unit of the pertinent source hemisphere that projected to the target unit, as defined by the outcome of that binomial trial, was determined in a subsequent step. This was done by random sampling from a uniform distribution where all units in that network-hemisphere have equal probability of being drawn.

#### 2.2.3 Density

The number of input connections each network unit receives from units in the preceding processing stage is controlled by a parameter, *d*, which we term density. This value is defined once for each instantiation of the model. This implies that all network units, by definition, receive the same number of input projections. For a density value of one, each unit in the target layer receives input from one source unit in the preceding layer. This unique source of input will determine its activation level to any given image. In contrast, for higher values of *d*, each unit in the target layer receives *d* projections from—most probably—multiple units in the previous layer. For density values larger than one, each unit integrates its multiple inputs by averaging the activation values of its source units. Network density values explored in the model were defined as 2^*q*^, with *q* = 0, 1, 2, 3, 4, 5.

### 2.3 Model variants

Two model variants were explored. In the first variant, termed pixel-level model variant (MV_pixel_), grayscale face images served as input to the network model (for example images, see Figure 4). Receptive fields of units in layers 1-8 were specified as explained in Section 2.2. Local luminance was the primary source of information propagated through this first variant of our model. In the second model variant, termed MV_S1_, instead of grayscale images, we relied on the S1 representation of the HMAX model as implemented by Serre et al. (2005). An advantage of this implementation over the original formulation by Riesenhuber and Poggio (1999) is that filter parameters (orientation, effective width, wavelength) were adjusted to match the tuning profiles of S1 units with those of V1 parafoveal simple cells. As in earlier versions of HMAX, the implementation by Serre et al. first applies a filter bank to each input image. Gabor filter banks of four orientations (0, π/4, π/2 and 3π/4 radians) and 16 receptive field (RF) scales (0.19 to 1.07 degrees of visual angle; 7 × 7 pixels to 39 × 39 pixels in steps of 2 pixels) thus lead to 64 filters, each associated with a unique pair of RF-size and orientation. All 64 filters are applied to each image location, and a measure hereby obtained of the energy matching the spatial frequency and orientation of each filter. In each image location, we summed filter output across all orientations and frequencies. We thus obtained a single intensity value per image location summarizing the total energy in the image matching the basis filters (for an example S1 filtered image, see Figure 4). This form of representation served as input to MV_S1_.

**Figure 4.**
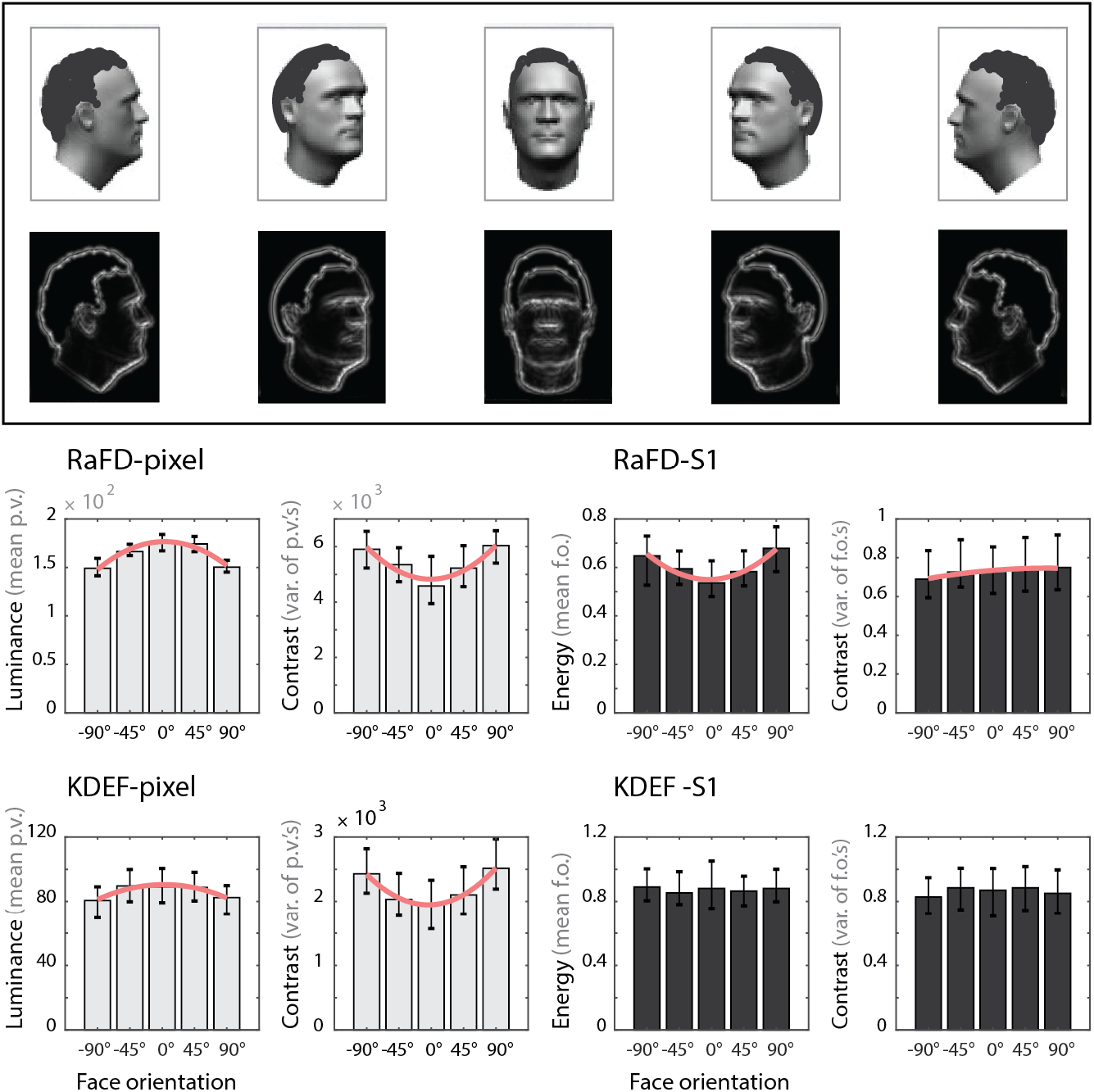
Distribution of mean luminance and contrast as a function of viewpoint for two face databases. Top row: example face identity shown in five orientations (90°, 45°, 0°, −45°, −90°). Second row: images above shown after pooling local orientation and frequency filters from the S1-layer of HMAX model. Below, median and interquartile range are shown for mean luminance (mean pixel value) or contrast (pixel variance) of face identities, always as a function of viewpoint. Bar plots in light gray correspond to pixel-level representation. Plots in black correspond to S1-level representation. Best-fitting second-order polynomial is shown in red only if either the linear or quadratic regression coefficients are significantly different from zero at the population level and their associated absolute values also significantly different. p.v. = pixel value; f.o. = filter output. Please note that all face images shown throughout this paper were computer generated and used for illustration purposes only, they are not instances of photographs from the analyzed databases.

The pixel-level representation captures biases in mean luminance across stimuli neglected by the S1-level representation. The S1-level representation was informed by electrophysiological measurements from macaque V1 and in this sense is more biologically plausible. Recent evidence revealing interactions between luminance and contrast responses in EVC (Vinke and Ling, 2020) suggest that these two representations provide complementary information and motivate their inclusion here.

#### 2.3.1 Event-related design

Images provided as input to the network were presented emulating an fMRI event-related design. Each stimulation event was defined by a unique combination of face identity and orientation. In the case of the RaFD database, this led to 57 (identities) × 5 (orientations) = 285 event types. In the case of the KDEF database, this led to 68 (identities) × 5 (orientations) = 340 event types.

### 2.4 Pattern generation and analyses

#### 2.4.1 Network activation patterns and their simulated measurement

In our network model, each unit models a single fMRI voxel. In turn, the collection conformed by all units in each hemisphere of a network layer define a region of interest (ROI)— e.g., layer 1, left hemisphere. Then, for each image provided as input to a specific instantiation of the network, patterns of activation are defined by concatenating the activation levels observed in each unit of the ROI under consideration. For example, response patterns to input images associated with a right hemisphere ROI in layer 1 would consist of 2048 entries, one entry per unit of the right layer 1 network hemisphere. In a similar fashion, the patterns associated with right hemisphere ROIs of layers 2-8 would each consist of 512 entries.

FMRI time series, one per voxel, naturally exhibit different levels of gain. Some fMRI voxels exhibit higher BOLD signal levels than others, due to, among many reasons, partial voluming (González Ballester et al., 2002), cortical folding (Polimeni et al., 2010; Gagnon et al., 2015) and specifics of the underlying vasculature (Bandettini and Wong, 1997; Schmid et al., 2019). A static measurement gain field (mGF) can be defined that summarizes the level of signal gain in each voxel (Ramírez et al., 2014; 2020). To model the influence of such gain field on our simulated response patterns, we randomly generated an mGF. For each network unit, we randomly sampled a scalar in the interval [0, 1] from a uniform distribution. Responses in each unit associated with each experimental condition were pointwise multiplied by its associated entry in the generated mGF, leading to simulated *measured* activation patterns in response to all images. We next used RSA to analyze patterns of activity across voxels within each ROI.

#### 2.4.2 Pattern similarity analyses

Representational Similarity Analysis (RSA) (Kriegeskorte et al., 2008) was used to analyze the simulated activation patterns. As in previous fMRI studies summarized in Table 1, our goal was to characterize the form of view-tuning of responses to face stimuli presented in various viewpoints. We conducted three RSA variants to investigate if, and if so, how, the form of the inferred viewpoint representations differed. These RSA variants differed with regards to two factors: (i) distance measure used to represent pattern dissimilarities (Euclidean, or correlation), and (ii) choice whether to demean the data prior to computing pattern dissimilarities.

The RSA procedure consisted of two main steps. In the first step, we computed empirical dissimilarity matrices (eDSMs) from the simulated activity patterns elicited by each face-identity in each region of interest. eDSMs were indexed by facial viewpoint (−90°, −45°, 0°, 45°, and 90°), leading to 5 by 5 dissimilarity matrices. Two distance functions were used to define eDSMs: the correlation distance (eDSM_corr_) and the Euclidean distance (eDSM_Euc_). Thus, two of our RSA variants were uniquely specified by the choice of pattern dissimilarity measure (i.e., RSA_corr_ and RSA_Euc_). The third RSA variant considered the same simulated activity patterns as before, but computed pattern dissimilarities after first demeaning the data. In the latter procedure, also previously referred to as “cocktail demeaning” (Haxby et al., 2001; Garrido et al., 2013), the mean response across conditions is subtracted from that observed for each condition in each measurement channel (e.g., each fMRI voxel). This operation has been implicitly and explicitly implemented in previous neuroimaging studies (see Table 1 and Ramírez, 2017). Since angular distances, but not Euclidean distances, are affected by data recentering, only RSA_corr_ was conducted on demeaned data. Thus, the option whether to demean the data defined the third RSA variant considered here—RSA_corrDem_.

In the second step, the computed eDSMs were compared with two representational models, each expressed as a model dissimilarity matrix (mDSM): (i) *Viewpoint model*, which assumes neuronal response patterns in some brain area reflect head angular disparity, and (ii) *Symmetry model*, which assumes neuronal populations respond similarly to mirror-symmetric face orientations, e.g., −90° and 90° views (see Figure 1). These model templates capture two forms of view-tuning observed in macaques (see Introduction). As measure of representational similarity (or, in other words, agreement between the rank-order of the entries in the model and empirical DSMs), the Spearman rank-order correlation was computed between eDSMs and mDSMs. These correlations only considered upper triangular matrix entries and excluded the main diagonal (Ritchie et al., 2017).

#### 2.4.3 Characterization of viewpoint representations at the single face-identity level

For each combination of face-identity and orientation, we calculated the Euclidean (i.e., L^2^) norm of each activation pattern from layer 8. We formed a *norm-profile* for each stimulus identity by concatenating the Euclidean norms of the patterns associated with each of the five face orientations. A general linear model (GLM) considering three regressors—viz., constant, linear, and quadratic—was fitted to each norm-profile using ordinary least squares (OLS) error minimization. The linear and quadratic regressors were mean-centered and normalized to unit length. The difference between the coefficient of determination (R^2^) of the linear and quadratic regressors was used to predict the degree of mirror-symmetry of the eDSMs associated with each face-identity. The linear association between these differences in R^2^ and the observed representational similarity with the Symmetry model was evaluated. A positive association would imply that the dominant form of bias observed on norm-profiles is predictive of the observation of mirror symmetry with RSA.

### 2.5 Image-level analyses

For whole-(see Figure 4) and half-image analyses (see Figure 5) we computed the mean and variance of the pixel- and S1-level representation of each face-identity presented in the five orientations considered in this study. Profiles of means and variances were formed as described in 2.4.3 for norm-profiles, but replacing the L^2^ norm of each activation pattern for the mean or variance of each associated face image. In the case of the pixel-level representation, the mean corresponds to mean image luminance, and the variance to image contrast. In the case of the S1-level representation, the mean corresponds to the mean energy matching the relevant Gabor filters (see Section 2.3 for details), and the variance corresponds to the variability of the pooled filter responses for the relevant input image. As performed above for norm-profiles, a GLM considering constant, linear and quadratic regressors was fitted to each mean- and variance-profile, and statistical tests conducted either on the regression coefficients, their associated R^2^, or the difference in R^2^ observed between the linear and quadratic regressors. The latter difference was used to characterize the dominant trend observed on these profiles—i.e., linear (antisymmetric), or quadratic (symmetric). We also explored a GLM additionally considering cubic and quartic regressors. Dominant trends were tested as above after adding the variance explained for even (quadratic and quartic) and odd (linear and cubic) polynomials. Here, we loosely refer to trends described by even polynomials as symmetric, and those described by odd polynomials as antisymmetric (but see Petitjean, 2021).

**Figure 5.**
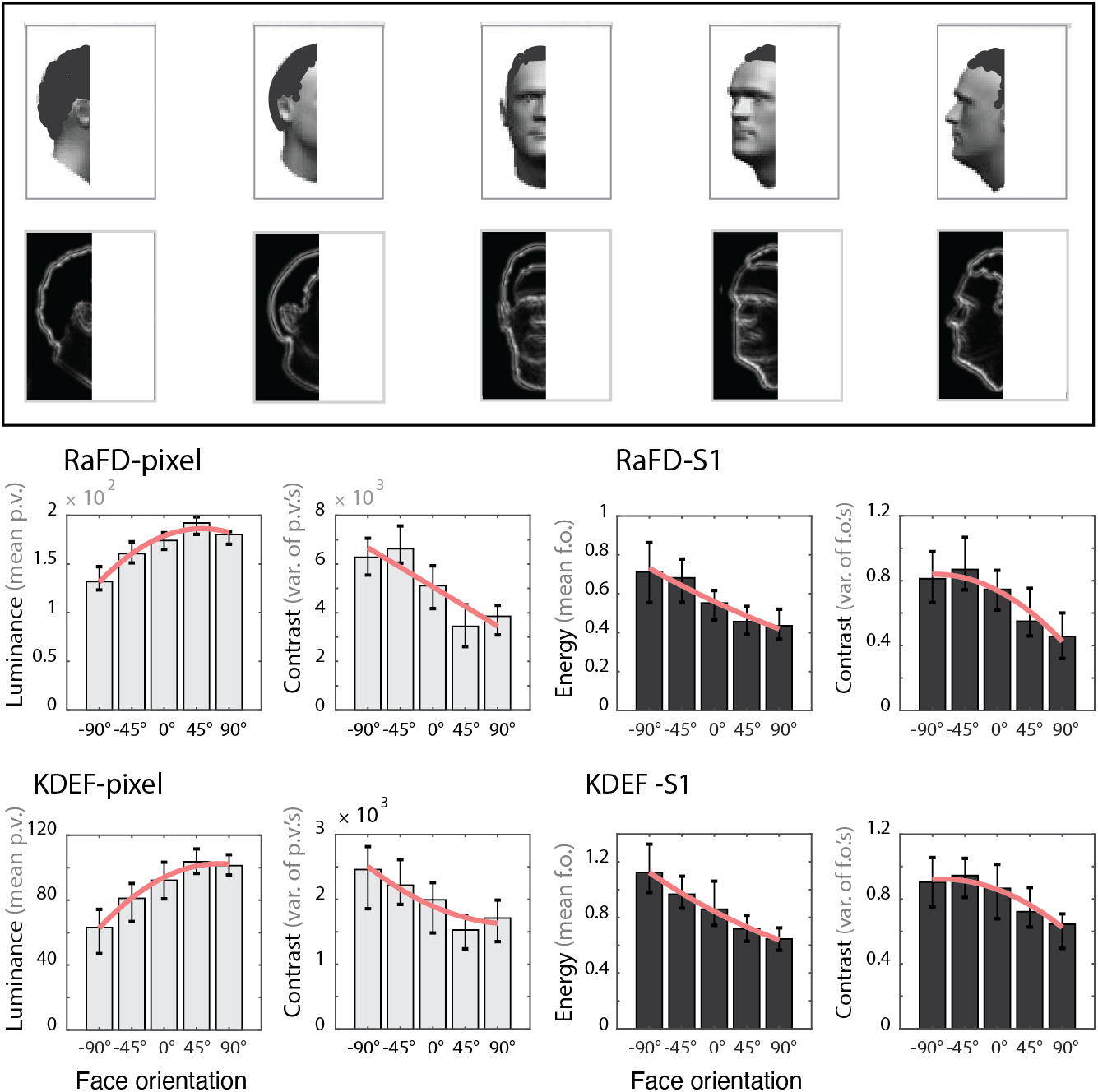
Distribution of mean luminance and contrast as a function of viewpoint for half-images for two face databases. Layout as in Figure 4. Here, however, only the left half of each image was analyzed. In contrast to Figure 4, where stronger quadratic than linear trends of mean luminance and contrast were usually observed across face views, linear trends proved dominant here regardless of database and representational format. Best-fitting second-order polynomials are shown in red following criteria in Figure 4.

### 2.6 Statistical analyses

For the sake of consistency, statistical tests reported throughout this paper were implemented by means of bootstrap tests on medians. All statistical analyses considered 10,000 bootstrap resamples. When testing in a population (i.e., face database) for systematic effects on GLM regression coefficients (see 2.4.3 for details), we tested the null hypothesis of zero median of the pertinent average regression coefficient by bootstrapping each database. All statistical tests were two-sided, unless a directional hypothesis is explicitly stated. To evaluate the dominance of linear over quadratic trends (and *vice versa*) irrespective of the sign of a polynomial trend across different face-identities in the population, we tested the null hypothesis of zero median across bootstrap resamples of the average difference of the R^2^ associated with the linear and quadratic regressors obtained for each face-identity. Similarly, to more generally test the dominance of symmetric over antisymmetric components, we tested the null hypothesis of zero median across bootstrap resamples of the average difference of the R^2^ associated with the even (square and quartic) vs. odd polynomial (linear and cubic) regressors.

Statistical tests for RSA were also implemented by bootstrap tests of medians. To evaluate a relative increment (or decrement) of viewpoint vs. mirror-symmetry when comparing early network layers (defined here as layers 1 and 2) and late layers (defined as layers 7 and 8), we compared in each combination of population (RaFD, KDEF) and model variant (MV_S1_, MV_pixel_) the difference in correlation between the simulated empirical DSMs and our two model DSMs (Symmetry, and Viewpoint). A relative increment (or decrement) when comparing early and late layers would indicate an interaction between hierarchical level of the network and the inferred prevalent form of view-tuning—e.g., a shift from a view-tuned representation in early layers to a predominantly mirror-symmetric representation in late layers. To test if the viewpoint and symmetry models each significantly correlate with the simulated data in early and late network layers, for each combination of population and model variant, we tested the null hypothesis of zero median of the Spearman correlation between the simulated data and the tested model. Finally, to compare the level of the network hierarchy at which the observed representational structure shifted from predominantly view-tuned to mirror-symmetric, we ran bootstrap tests on medians of the average zero-crossing of the difference between the viewpoint and mirror-symmetric models.

## 3. RESULTS

Of seven studies investigating viewpoint representations in humans using fMRI-MVPA, five focused on brain activation patterns found to be reliable within individual subjects (see Table 1). Our model primarily addresses these five studies. The remaining two studies relied on across-subject RSA approaches unsuited to draw inferences about spatially-structured brain patterns reliable at the single-subject level (Sabuncu et al., 2010; Haxby et al., 2011; Yamada et al., 2015; Feilong et al., 2018). We have previously addressed the interpretation of these types of studies (Ramírez et al., 2020). Of the five within-subject fMRI-MVPA studies in Table 1, three considered a task that required subjects to fixate on a known image location (i.e., Axelrod and Yovel, 2012, Experiment 2; Kietzmann et al., 2012; Ramírez et al., 2014), and for this reason are the prime focus of our model. Axelrod and Yovel (2012, Experiment 1) relied instead on a one-back identity task that allowed subjects to freely move their eyes. If the antisymmetric pattern of activations across face-views reported in EVC, attributed by the authors to systematic eye-movements towards the facial features (e.g., eyes and mouth), is assumed to generalize to the study by Guntupalli et al. (2017), which relied on a similar task, our model provides a straightforward account of these two additional experiments. The central intuition behind the account proposed here is that, given previously observed low-level image biases across face-views, considering a system exhibiting cortical magnification and a gradually increasing degree of interhemispheric connectivity is sufficient to account for key common and inconsistent trends observed among within-subject studies.

We present our results in four steps. First, we report analyses demonstrating the prevalence in two popular face databases of biases in the distribution of low-level features (i.e., mean luminance and contrast) across face-views qualitatively like those previously noted in a smaller set of face-stimuli (Ramírez et al., 2020). Second, we present simulation results probing the impact of cortical magnification (CM) on the manifestation of these low-level biases in the input layer of our network model. Third, we present RSA results of response patterns to face-images obtained from our network model. We focus on the impact of two analysis choices—i.e., pattern dissimilarity measure, and data recentering—on the inferred form of the underlying viewpoint representations: view-tuned, or mirror-symmetric (see Figure 1). Finally, we probe the efficacy of a simple statistic based on the observed signal-strength of the simulated response patterns to predict the degree of mirror-symmetry observed in ensuing pattern analyses at the single face-identity level.

### 3.1 Image-level analyses: whole- and half-images

To test whether low-level feature imbalances of the type assumed by our model are prevalent among face stimuli, we investigated the distribution of mean-luminance and contrast across facial viewpoints (−90°, −45°, 0°, 45°, 90°) of two popular face databases—the KDEF and RaFD databases. Figure 4 presents results for analyses of full images. The goal of these analyses was to uncover biases across conditions expected to manifest in brain areas populated by neurons with RFs that cross the vertical meridian into the ipsilateral hemifield. Two representational formats of images from these two databases were investigated. The first format was the pixel-level representation. The second format focused on the pooled output of frequency- and orientation-tuned filters of the S1 layer of HMAX in the biologically constrained implementation used by Serre et al. (2005) to model receptive fields of V1 simple cells (see Methods). We term this second format S1-level representation.

Pixel-level analyses revealed systematic quadratic biases about the frontal face-view in both databases (Figure 4). Such biases were evident for both mean luminance and contrast. Statistical tests confirmed that when each database was treated as a random sample of face identities from some population, quadratic trends on regression coefficients were significantly different from zero (bootstrap test on medians, all *p* < 0.001). For both analysis variants and databases, quadratic trends were found to be significantly stronger than their linear counterparts in terms of R^2^ (bootstrap test on medians, all *p* < 0.001). As can be observed in Figure 4, the sign of these quadratic biases was congruent across databases.

However, for S1-level analyses only the RaFD database revealed a significant quadratic trend, specifically, for the distribution of mean filter energy observed across facial viewpoints (bootstrap test on median: *p* < 0.01) (Figure 4). Note that this analysis is based on averages of S1 filter responses instead of pixel values, and the observed trends therefore not necessarily proportional to mean luminance. These two measures provide distinct information about the images. When analyzing the variance of the S1 filter energy observed across image locations (an analysis similar to that of image contrast across views but, again, now based on S1 filter outputs instead of pixel values), only the linear regression coefficient for the RaFD database proved significantly different from zero (bootstrap test on median: *p* < 0.001)). Further analyses revealed that the magnitude of the median regression coefficient for the quadratic regressor was, nonetheless, larger than its linear counterpart, and this difference statistically significant (bootstrap test on median, *p* < 0.001). This result is explained by larger variability among the quadratic than the linear regression coefficients. Crucially, similar analyses at the level of R^2^ (instead of regression coefficients), which disregard the sign of the analyzed polynomial trends for each face identity, revealed significantly stronger quadratic than linear trends regardless of low-level feature, database, and representational format (bootstrap test on medians: all *p* < 0.001). In sum, analyses on whole images revealed significant mirror-symmetric biases across face-views. Critically, quadratic components always significantly outweighed their linear counterparts in terms of R^2^. These results show that mirror-symmetric low-level biases about the frontal view are common among face stimuli, in line with previous observations (Ramírez et al., 2020). We next asked if the opposite pattern of results is observed for half-images, as assumed by our model (see Figure 2). We analyzed half-images to see if the linear trend in this case outweighed the quadratic trend.

Half-image analyses otherwise identical to those reported for full-images in Figure 4 are summarized in Figure 5. These analyses aim to characterize low-level biases across face-views due to mean-luminance and contrast. Such biases are likely to be observed in areas with RFs circumscribed to one visual hemifield, such as V1. As hypothesized, a clear dominance of the linear over the quadratic trend was observed for both low-level features, regardless of database, when analyzing regression coefficients (bootstrap test on medians, all *p* < 0.001). A similar pattern of results was observed for the mean and variance of the S1 filter outputs (bootstrap test on medians, all *p* < 0.001). For completeness, we decomposed the profiles for each face-identity into linear and quadratic components, as done for analyses of the full images (see Figure 4). We found that the linear component was significantly stronger than its quadratic counterpart at the level of R^2^ regardless of low-level feature, database, and representational format (bootstrap test on medians, all *p* < 0.001). This implies a double dissociation for full and half-images with regards to their dominant polynomial trends—namely, quadratic over linear for full images, and linear over quadratic for half-images. In sum, we found evidence of low-level biases in half-images consistent with the antisymmetric biases assumed by our model, as well as evidence of the dominant symmetric biases about the frontal face-view anticipated for full images.

### 3.2 Impact of cortical magnification and network density on Layer 1 activation profiles

If biases across face views like those described in Figures 4 and 5 were present, respectively, in FFA and V1, this would suggest a simple account of key similarities and inconsistencies observed across the fMRI-MVPA studies summarized in Table 1 (see also Figure 1). However, it may be argued that image-level analyses inadequately represent biases that exist at the level of images when it comes to consider their influence on V1 activation profiles. Crucially, both image-level representations considered here lack a fundamental property of primate V1, namely, cortical magnification (CM). To assess whether biases qualitatively like those reported in Figures 4-5 are observed in a representation considering CM, we conducted analyses homologous to those reported in Figures 4-5, but now on activation patterns associated with the same images in Layer 1 of the feedforward randomly-connected two-hemisphere network proposed here (see Figure 3).

First, we assessed whether the mean activation observed across all Layer 1 units (roughly analogous to analyzing whole images) or, alternatively, across only one hemisphere of Layer 1 (roughly analogous to analyzing half-images) exhibit trends qualitatively like those found at the pixel- and S1-level. When jointly considering units from both hemispheres of Layer 1, as in full-image results, we again found marked mirror-symmetric biases as a function of viewpoint for both mean and variance of Layer 1 activation patterns. However, we also noted that the *positive* quadratic trend across views previously observed for contrast (pixel-level, whole image) now exhibited a significant *negative* quadratic trend (see Figure 6*a*, top). In a similar vein, we noted that the dominant linear trend found for contrast for RaFD-S1 changed to positive quadratic (see Fig. 6*a*, bottom). These observations demonstrate that although cortical magnification does influence the precise form and strength of the mirror-symmetric biases noted when analyzing image-level representations, marked mirror-symmetric biases of the form assumed by our model are again observed when probing Layer 1 activation profiles. In turn, analyses of patterns from only one hemisphere of Layer 1 revealed dominant antisymmetric biases consistent with those shown in Figure 5 for half-images.

**Figure 6.**
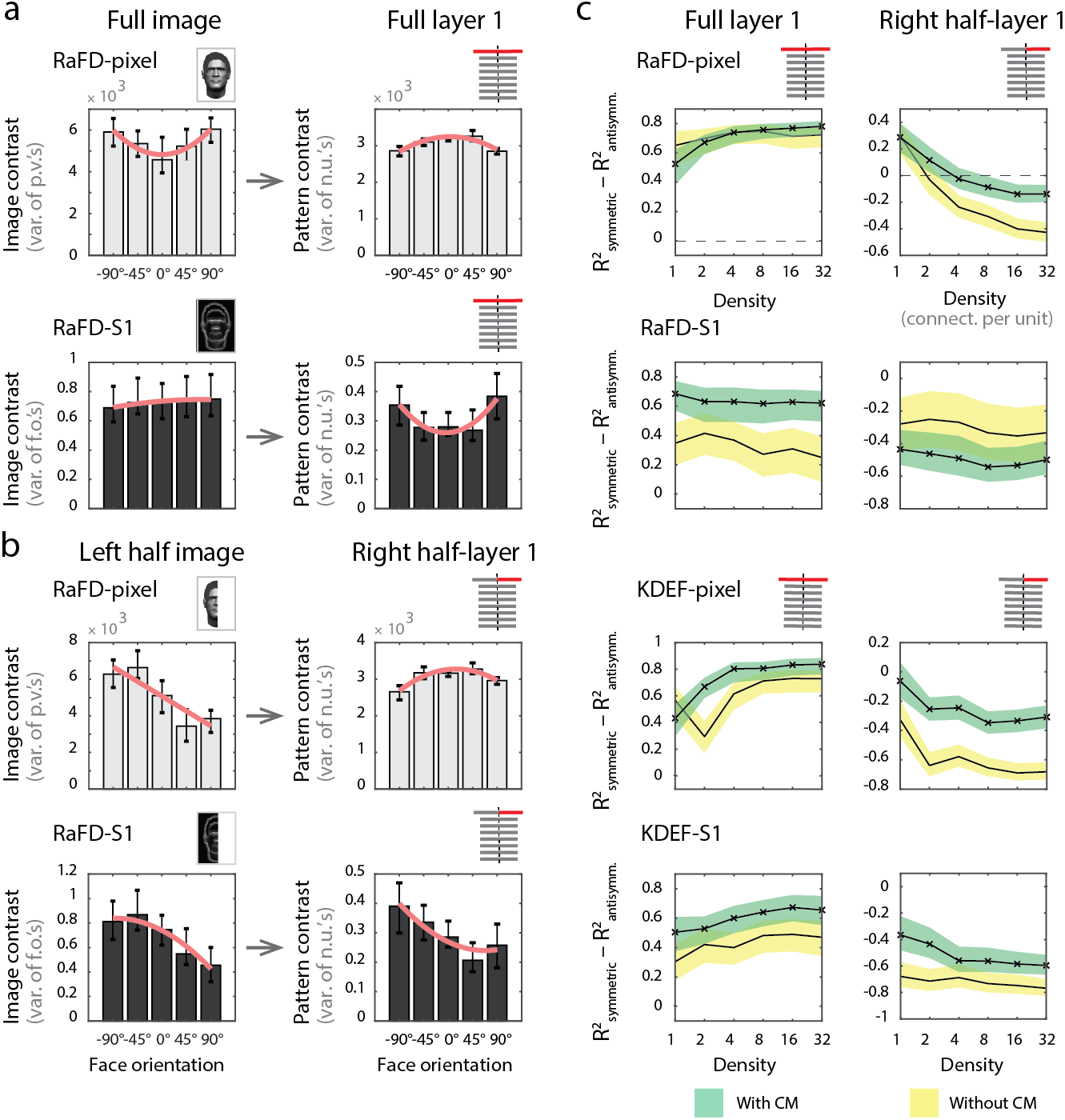
Impact of cortical magnification and network density on Layer 1 activation profiles. *a-b*) Results are organized according to portion of the image (whole, or left-half) or network hemispheres analyzed (both, only right). Image- and network-level analyses are shown, respectively, on the left and right column. Analyses computed on pixel- and S1-level representations are shown, respectively, on the top and bottom row. *a*) Top, left: Median image contrast across face identities for each viewpoint. Error bars indicate interquartile ranges. Best fitting second order polynomials are overlayed on each plot. Top, right: Median variance of activation patterns associated with each face identity in Layer 1. Unlike image-level analyses shown to the left, network analyses shown to the right consider cortical magnification of central image locations. *b*) Median contrast across face identities as a function of viewpoint for half-images. Panel is organized as panel *a*. Note differences in form and direction of trends when contrasting image- and network-level analyses. *c)* Difference in R^2^ of symmetric (quadratic and quartic) and antisymmetric (linear and cubic) trends for Layer 1 activation patterns as a function of network density (x-axis) and cortical magnification (green = CM, yellow = no CM). As in panel *a*, activation profiles for each face identity were formed by concatenating the variance of activation patterns for each face view. Shaded areas indicate 95% confidence intervals. Positive values indicate stronger symmetric than antisymmetric trends. Note consistency in the direction of dominant trends regardless of database, cortical magnification, and number of hemispheres. p.v. = pixel value; f.o. = filter output; n.u. = network unit.

Next, we evaluated if the pattern of results in Layer 1 of our network model is specific to the lowest density level, as reported in Figure 6*a*-*b*, or prevalent over a wider range of densities. High sensitivity to network density would in our view challenge our model as a plausible explanation of empirical data. In contrast, robustness over a wider range of densities would lend plausibility to the model. To address this issue, we quantified the relative dominance of symmetric and antisymmetric trends across face views in each network layer as a function of network density. Profiles describing the signal strength across face-views for each face identity were specified by concatenating the variance of the activation pattern associated with each face view for that identity. The relative strength of symmetric (even) and antisymmetric (odd) polynomial trends on these activation profiles was evaluated. We did this for both databases (RaFD and KDEF) and representational formats (pixel and S1). To assess the impact of cortical magnification on network behavior, we also ran a set of simulations ignoring this property. As observed in Figure 6*c*, while cortical magnification and network density were found to modulate the relative influence of the analyzed symmetric and antisymmetric trends, critically, the direction of these biases was consistent across network density levels. The only exception was the observation for the lowest density level (*d*=1) of a dominant symmetric trend for half-layer network analyses for MV_pixel_ for the RaFD database (*see* Figure 6*c*, top row, right column). Except for this observation (see Discussion), as expected, a predominance of the antisymmetric component was evident for densities larger than two regardless of image database and model variant. Symmetric and antisymmetric biases of the form assumed by our model were found with and without cortical magnification.

### 3.3 RSA analyses of simulated activity patterns

The biases described thus far are agnostic regarding the spatial structure of the underlying distributed response patterns. The reported analyses are only informative regarding the mean and variance of the simulated brain patterns as a function of viewpoint. According to our account, such biases may be sufficient to explain the dominant trends observed across the fMRI-MVPA studies considered here. Naturally, we next asked if analyses of the patterns associated with face images of the two datasets reveal biases sufficient to account for the consistencies and inconsistencies summarized in Figure 1.

To address this question, we subjected activation patterns in each network-layer to Representational Similarity Analyses (RSA) (see Methods for details). We computed (simulated) empirical dissimilarity matrices (eDSMs) describing the dissimilarity relationships of the five facial viewpoints for each face identity. eDSMs were computed according to two distance measures—correlation distance, and Euclidean distance. Because angular relationships across pattern-vectors are expected to change after data recentering (e.g., after mean-centering) only with angular, but not Euclidean distances, we also report RSA on demeaned data when using the correlation distance to measure pattern dissimilarities. For analyses on demeaned data, the mean response across experimental conditions was computed for each network unit and subtracted from the value associated with each condition, as previously done for fMRI voxels (see Table 1). We argue that these three analysis strategies (RSA_corr_, RSA_Euc_, and RSA_corrDem_) capture key analysis choices across studies that may largely account for the observed inconsistencies. Pattern analysis results are summarized in Figures 7 and 8. We first report simulations considering a network density of 16 connections per unit. As expected, analyses relying on the correlation distance led to the observation of markedly view-tuned representations throughout the eight network layers (Figure 7, left). For both face databases and model variants, the monotonically view-tuned model exhibited higher correlations with the simulated eDSMs than the mirror-symmetric model. The only exception was for the S1 model-variant (MV_S1_) for the RaFD database, which exhibited in Layer 1 higher correlations with the mirror-symmetric than the view-tuned model (see Discussion).

**Figure 7.**
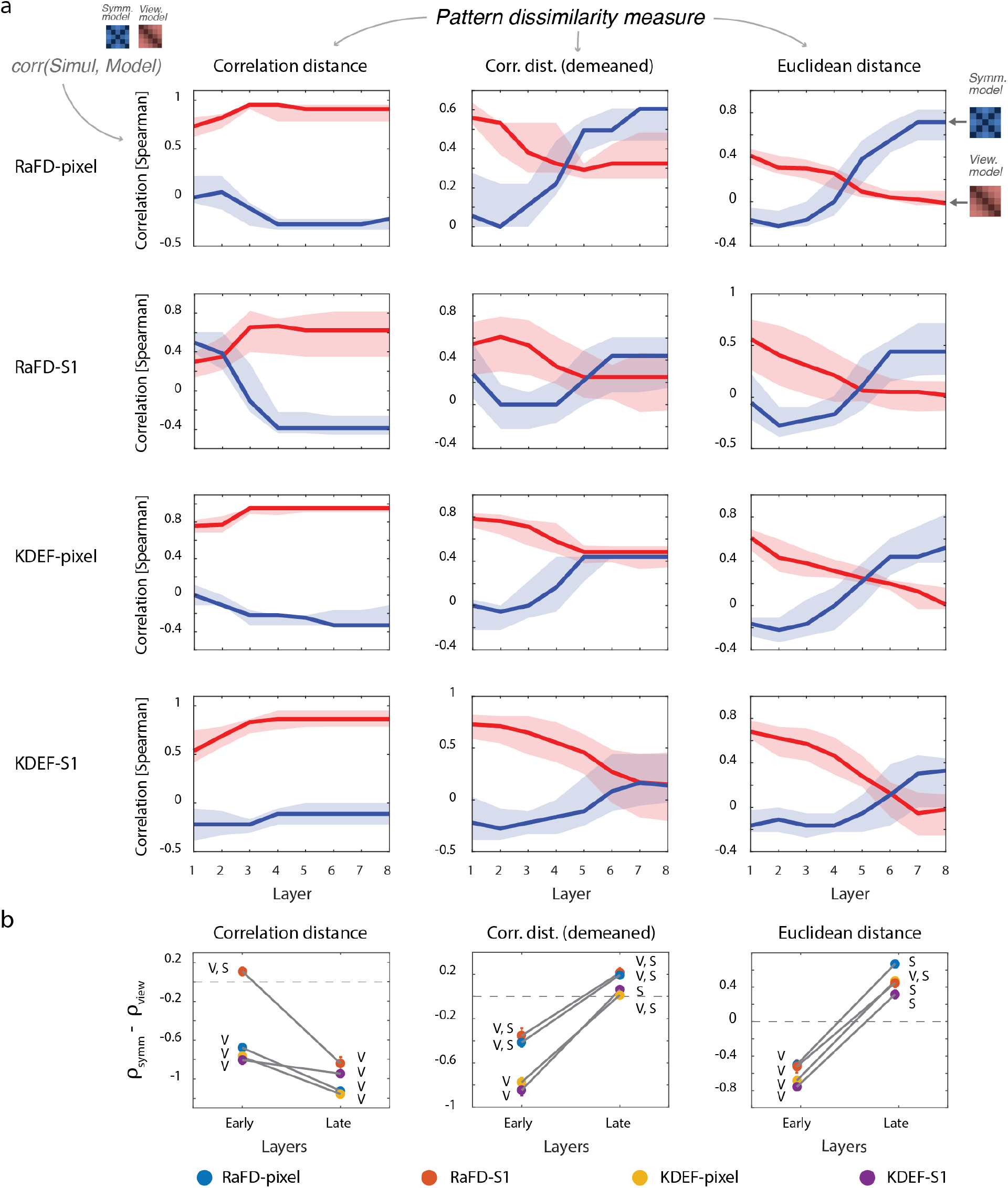
RSA results across network layers for density-level = 16. *a*) Each sub-plot summarizes RSA outcomes for network-layers 1-8 for a single combination of image database, model variant, and RSA variant. Rows indicate image database (KDEF, RaFD) and model variant (pixel, S1) combinations. Columns indicate RSA variant: correlation distance (RSA_corr_), correlation distance on demeaned data (RSA_corrDem_), and Euclidean distance (RSA_Euc_). For each network layer (x-axis), median correlations are shown between simulated empirical dissimilarity matrices and the view-tuned model (in red) and the mirror-symmetric model (in blue). Shaded areas indicate interquartile ranges. Note consistently dominant view-tuned representations across layers for RSA_corr_ (except for RaFD-S1 at layer 1). In contrast, however, for RSA_corrDem_ and RSA_Euc_ a relative decrease of view-tuning and increase of mirror-symmetry is observed along the hierarchy. *b*) Average correlation difference between the symmetry and viewpoint models (y-axis), in early (1 and 2) and late (7 and 8) network layers. “V” and “S” indicate that viewpoint and symmetry correlations, respectively, are significantly above zero. While RSA_corr_ exhibits significantly higher view-tuned representations across layers, RSA_corrDem_ and RSA_Euc_ exhibit a clear shift from viewpoint to mirror-symmetry, or a mixed representation, when comparing early and late layers. Gray lines indicate statistically significant increments or decrements when comparing early and late network layers (bootstrap test of medians, *p* < 0.001). For RSA_corrDem_ and RSA_Euc_, significant increments in mirror-symmetry are observed in late layers compared with early layers for all model variants and image databases. For RSA_corr_ an increase of view-tuning is observed instead.

**Figure 8.**
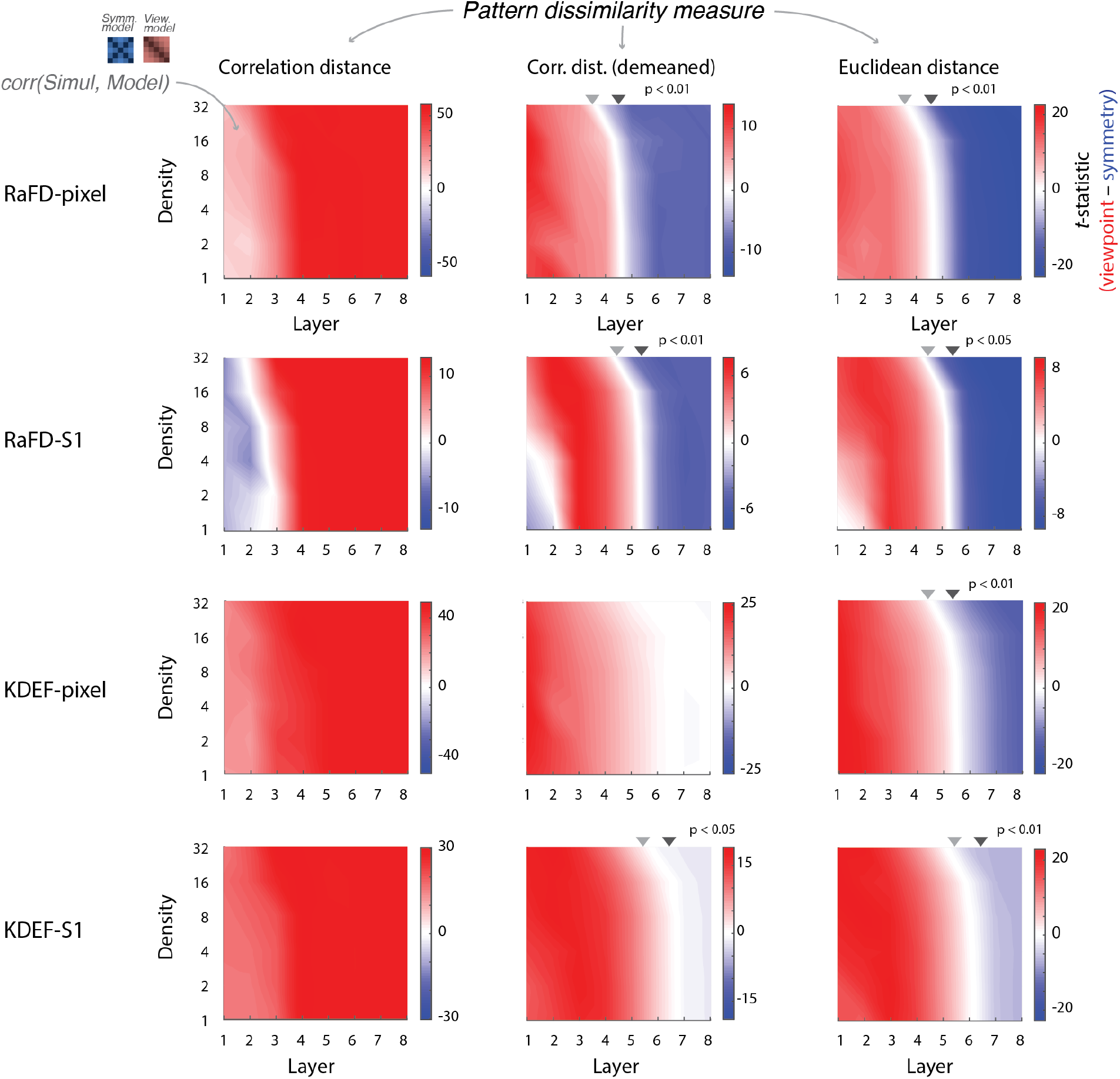
RSA results for multiple network densities. Layout as in Figure 7. Here, however, y-axis denotes density-level of the analyzed patterns. Each plot color-codes the paired *t*-statistic comparing correlation coefficients of the simulated empirical DSMs with the Viewpoint and Symmetry models. Areas in red indicate consistently higher correlations with the viewpoint model. Areas in blue indicate consistently higher correlations with the Symmetry model. Results are broadly concordant with those shown in Figure 7 for a network density of 16 (see text for one exception). Dark-gray and light-gray triangles, respectively, indicate for the lowest and highest network-densities (*d* = 1 and 32) the point at which a shift occurs from a predominantly view-tuned to a predominantly mirror-symmetric representation. Triangles are only shown when the difference in the point of zero-crossing for the two network densities was statistically significant. In 3/4 RSA_corrDem_ and 4/4 RSA_Euc_ analyses, this shift occurs earlier for the highest than the lowest density level.

Analyses relying on the correlation distance but—crucially—now conducted on demeaned data, as well as analyses relying on the Euclidean distance, led to marked incremets in later network layers in the association between the simulated empirical DSMs and the mirror-symmetric model (see Figure 7*a*). As further shown in Figure 7*b*, correlations between the mirror-symmetric mDSM and the simulated data were found to be significantly larger than zero, as well as to increase significantly in later layers (defined as 7 and 8) with respect to earlier layers (defined as 1 and 2) (bootstrap tests on medians: all *p* < 0.001). Moreover, in 2/4 combinations of database and model variant for RSA_corrDem_, and in 4/4 combinations for RSA_Euc_, the mirror-symmetric model significantly outperformed the view-tuned model in later layers (Figure 7*b*). For RSA_Euc_ and RSA_corrDem_ no instance was found where the view-tuned model significantly outperformed the mirror-symmetric model. In sum, these observations show that a parsimonious model incorporating cortical magnification and gradually increasing interhemispheric crossings, given the observed low-level image biases, leads to geometric configurations in multivariate pattern space that account for inconsistencies observed in higher-level processing stages such as FFA. Specifically, these inconsistencies arise due to the influence on RSA outcomes of data analysis choices—namely, pattern dissimilarity measure and whether or not to demean the data prior to RSA. Our observations also explain the view-tuned pattern of response consistently reported in EVC (see Table 1).

Next, we assessed whether the pattern of results observed for density 16 is consistent across a broader range of network densities. Identical analyses as those reported in Figure 7, but for a wider range of densities, are summarized in Figure 8. These analyses show a broadly consistent pattern of results within the range of network densities explored. However, we observed two unexpected interactions. First, RSA_corrDem_ for the S1-model variant for the RaFD database revealed an interaction between network layer, density, and the observed representational structure. In more detail, only for early network layers (1 and 2) and low densities, the mirror-symmetric model outperformed the viewpoint model (see Figure 8, second row, middle column). In contrast, when the data were *not* demeaned, the mirror symmetric model outperformed the viewpoint model in these same layers regardless of network density (see Figure 8, second row, left column). Second, the network layer at which the observed representations changed from predominantly view-tuned to mirror-symmetric for 4/4 RSA_Euc_ and 3/4 RSA_corrDem_ analyses occurred slightly earlier for the highest density (*d* = 32) than the remaining densities. Statistical analyses showed that these differences are significant in the two populations (bootstrap tests on medians, all *p* < 0.05). These observations indicate that network density can significantly influence the outcome of RSA results. Overall, and as expected, analyses reported in Figure 8 revealed view-tuning with RSA_corr_ regardless of network layer, and RSA_corrDem_ and RSA_Euc_ led to the observation of view-tuning in early layers and mirror-symmetry in later layers. Crucially, data demeaning induced mirror-symmetry in the simulated eDSMs only in later network layers (see Figures 7*b* and 8). By the contrary, data demeaning induced view-tuning in early network layers. This can be observed in Figure 8 when comparing the color of higher network-densities in Layer 1 for RSA_corr_ and RSA_corrDem_ for the RaFD-S1 model; while RSA_corr_ looks blue (i.e., mirror-symmetric), RSA_corrDem_ looks red (i.e., view-tuned). See Figure S1 for supplementary analyses and discussion.

### 3.4 Signal-strength across face-views: Linear and quadratic trends predict RSA outcomes

Thus far, we found that overall trends in RSA analyses probing for viewpoint and mirror-symmetric representations on network-level analyses consistently confirmed biases of the form we initially hypothesized (see Figure 2). We noted, however, substantial variability across face identities for the outcome of RSA analyses (see Figure 7*a*). If our assumption is correct that the difference in R^2^ between quadratic and linear components computed on the profiles describing the strength of the signal observed for each face-view determines the outcome of RSA results, then, this difference should predict the degree of mirror-symmetry observed in Layer 8 with RSA at the single face-identity level. Figure 9 summarizes the results of the suggested analysis for the S1 variant of our network model. As hypothesized, the difference in R^2^ between quadratic and linear components was significantly positively correlated with the degree of mirror-symmetry observed with RSA_Euc_ and RSA_corrDem_ (range of *r* values: [0.69, 0.81], all *p* < 0.001), but not with RSA_corr_ (KDEF: *r* = −0.19, *n.s*., RaFD: *r* = −0.56, *p* < 0.01). Qualitatively similar results for pixel-level analyses are shown in Figure S2 and with an alternative norm-profile decomposition in Figure S3. These results show that biases in signal strength across views at the single face-identity level are predictive of the degree of mirror-symmetry observed with RSA in the last layer of our model.

**Figure 9.**
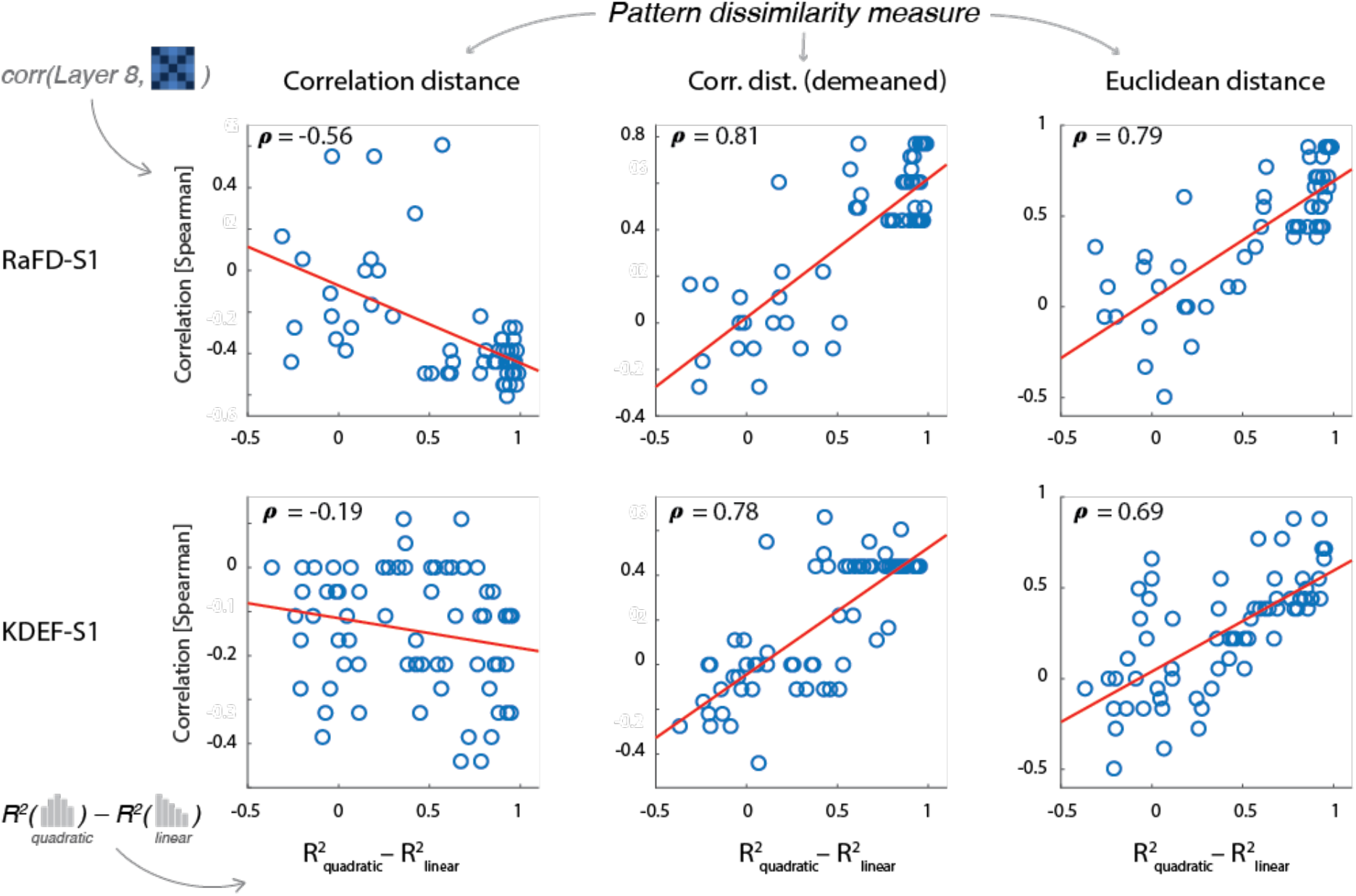
Accounting for mirror-symmetry in RSA for single face identities. Columns are indexed by pattern dissimilarity measure of the RSA variant used to analyze response patterns from Layer 8 of network model. Rows are indexed by image database. Points in scatterplots represent specific face identities. For each viewpoint (−90° to 90°, step 45°) of each face identity, we calculated the length (L^2^ norm) of its associated activation pattern. For each face identity, concatenating the norms for its five viewpoints served to define a norm-profile. A general linear model with constant, linear, and quadratic regressors was fit to each norm-profile. The difference in R^2^ between the quadratic and linear regressors (x-axis) is plotted against the degree of mirror-symmetry observed with RSA for each face identity (y-axis). Best fitting regression lines are shown in red. Outcomes of RSA_Euc_ and RSA_corrDem_, but not RSA_corr_, positively correlate with the degree of mirror-symmetry observed for single face-identities (all *p* < 0.001).

## 4. DISCUSSION

This study evaluated the ability of a parsimonious model to explain concordant and discordant trends observed across fMRI-MVPA studies investigating facial viewpoint representations in human visual cortex. These include reports of view-tuning in early visual cortex (EVC), as well as, in some cases, a gradual increase in the degree of mirror-symmetry as one proceeds along the visual hierarchy. Outstanding inconsistencies regard the very report of mirror-symmetry in mid- and high-level visual areas. Our hypothesis was that a simple biologically-inspired model incorporating three constraints—convergent feed-forward projections, gradually increasing interhemispheric connections, and cortical magnification— might reconcile these observations. Our simulations considered the influence of a measurement gain field, capturing a key and often-overlooked aspect of fMRI measurements. Our simulations explain why conclusions in EVC are consistent across studies regardless of data recentering and pattern dissimilarity measure, while conclusions in higher-tier areas depend on these choices. Our results also suggest that fMRI-MVPA studies may have misinterpreted low-level feature imbalances across conditions as evidence of mirror-symmetrically tuned neuronal populations. Below we discuss our findings, strengths and limitations of our model, and the broader significance of our findings.

### Low-level biases: linear and quadratic trends as a function of viewpoint

Image-level analyses of two popular databases replicated previous observations of mirror-symmetric biases about the frontal face view (Ramírez et al., 2020). Analyses of half-images instead revealed antisymmetric biases. To test whether such biases are modulated by cortical magnification, we probed layer 1 of our network model. While the sign and strength of trends across views were modulated by CM, they remained significant. Our simulations show the influence of low-level biases cannot be evaluated by simply inspecting pixel-level or wavelet-pyramid representations of the stimuli. They demonstrate CM should therefore be considered in models aiming to distinguish the impact of low-level features in visual cortex from other variables of interest, such as category structure and form of view-tuning.

### Simulation results

RSA of simulated patterns associated with face-images from both databases replicated key empirical observations. As observed in FFA, we found mirror-symmetry in later network layers with RSA_Euc_, but view-tuning instead with RSA_corr_. This is consistent with observations using classification-rates from multiclass linear support vector machines (SVM) as proxy for pattern dissimilarity (Axelrod and Yovel, 2012). The latter metric is sensitive to the Euclidean distance between pattern-vectors. In turn, relying on RSA_corr_, another study reported evidence of view-tuning in right FFA, and no mirror-symmetry (Ramírez et al., 2014). Both studies, however, observed stronger responses for frontal-than lateral face-views in FFA, as indicated by significant negative-quadratic trends in univariate analyses. Such trends are expected to lead to the observation of mirror-symmetry with RSA_Euc_. Indeed, Ramírez et al. observed partial mirror-symmetry in FFA with RSA_Euc_, but no evidence of mirror-symmetry with RSA_corr_. These observations are consistent with response biases revealed by our model and offer a unitary explanation for the inconsistent conclusions reached in these two studies.

A geometric principle behind our model is that data recentering does not change Euclidean distances among fMRI pattern-vectors, but changes their angular distances. Moreover, mean-centering can induce mirror-symmetry on eDSMs if mirror-symmetric biases in signal-strength are present in the data (Ramírez, 2017). Our simulation results support these concepts and help explain a reported gradual increase of mirror-symmetry, first noticed in OFA, but found to be prevalent both in the ventral and dorsal streams with an RSA_corrDem_ searchlight approach (Kietzmann et al., 2012). This finding is reminiscent of a reported gradual increase in BOLD sensitivity to ipsilateral stimuli along the ventral stream (Hemond et al., 2007). If data exhibiting a symmetric bias that increasingly manifests along the visual hierarchy is mean-subtracted, our model predicts, widespread symmetry beyond V1/V2 should gradually emerge and maximally manifest at a stage analogous to FFA. Another fMRI study found no evidence of mirror-symmetry in visual cortex, but widespread view-tuning, with a searchlight approach based on linear SVM classifiers (Guntupalli et al., 2017). The absence of a fixation task in this study, together with pronounced linear univariate trends across views observed in EVC with a similar task (Axelrod and Yovel, 2012, Exp. 1) may account for this difference (for details, see Supplementary Discussion).

All studies detailed in Table 1 observed view-tuning in EVC. Our model reproduced these observations regardless of database and model variant—one exception is discussed further below. View-tuning is observed in early network layers because the length of pattern vectors associated with different face-views, as well as their associated angular distances, are both antisymmetric. In other words, both tend to increase monotonically with head angular disparity. Hence, both RSA_corr_ and RSA_Euc_ lead to view-tuned eDSMs, since the former is sensitive to pattern-vector angles and the latter sensitive to their lengths. Mean-subtraction in the presence of anti-symmetric biases on vector norms will induce anti-symmetry in RSA analyses with angular distances, and eDSMs will hence remain antisymmetric (i.e., “view-tuned”) after demeaning the data. Our model thus explains findings of view-tuning in EVC regardless of analysis choices. In contrast, RSA outcomes in later network layers depend on distance measure and data recentering because of the inconsistent trends observed on vector norms and angles—mirror-symmetry on vector norms, antisymmetry on angles. Mean-centering across conditions in the presence of a quadratic trend on vector norms will induce mirror-symmetry on ensuing RSA analyses with angular distances. Thus, the low-level confounds described in Figure 5 imply a bias to observe view-tuning in EVC even if face-images happened to exhibit mirror-symmetric biases in their spatial structure—as has been previously suggested (Kietzmann et al., 2012). In turn, confounds described in Figure 4 imply a bias to observe mirror-symmetry in higher-tier visual areas with RSA_Euc_ and RSA_corrDem_, ironically, even if the spatial structure of the images happened to exhibit antisymmetry.

Unlike empirical observations in EVC, one simulation revealed mirror-symmetry in early network layers: S1 model-variant, RaFD database. This bias was observed with RSA_corr_ and RSA_corrDem_ (only for lower densities), but not RSA_Euc_. This finding reflects a genuine mirror-symmetric bias in the spatial structure of images from this database. The interaction between inferred form of view-tuning, network-density, and RSA distance measure is explained by the logic introduced above; mean subtraction in the presence of an antisymmetric bias among patter-vector norms induces biases on RSA consistent with the view-tuned model template. Mean-centering overly influenced RSA outcomes for early network-layers and higher densities because the strength of antisymmetric biases on vector norms was proportional to network density (see Supplementary Figure S1). Observations at higher network densities—presumably more comparable to fMRI measurements—and simulations sensitive to mean luminance (i.e. MV_pixel_) did not reveal this mirror-symmetric bias. In sum, the observed biases are broadly compatible with the empirical observations summarized in Table 1.

### Model strengths and limitations

Strengths of our model include its simplicity, ability to explain a range of observations, biologically-motivated constraints, and generalization across a range of network-densities and two databases. However, the primate brain exhibits feedback connections (Salin and Bullier, 1995; Briggs, 2020), category structure (Op de Beeck et al., 2019), lateralization (Rossion, 2014), and neuronal tuning properties that support behavior (Hubel and Wiesel, 1962; Gross et al., 1972; DiCarlo et al., 2012; Afraz et al., 2015). Our model does not incorporate these properties. Ours is not a realistic model of the visual system, but a biologically-informed device to comprehend the behavior of pattern analyses under minimal assumptions. The model is not directly informative regarding the form-of-tuning of neurons in any visual area. The model does reveal, however, sources of variance that must be considered to draw valid inferences regarding neural population codes with fMRI-MVPA.

An implicit assumption of our model is that facial viewpoint representations in face-selective areas do not markedly depend on task. A recent study supports this notion (Foster et al., 2022). Our model indicates, however, that eye-movements can strongly impact fMRI-MVPA observations. For this reason, our model mainly addresses studies where subjects maintained fixation at the center of the screen. Information about eye position, or methods to extract it from BOLD fMRI measurements (Frey, Nau, et al., 2021), may help extend our model.

### Overrepresentation of the frontal face views

The influence of low-level features in visual cortex is well-documented (Andrews et al., 2015). Yue et al. (2011) found that, on average, fMRI responses in EVC and FFA were stronger for frontal than lateral face views. From the strong correlation observed between EVC and FFA responses, they argued the latter may be caused by the former. Several studies have observed an overrepresentation of the frontal face-views in ventral visual cortex (reviewed in Ramírez, 2018). The question remains whether this overrepresentation is due to low-level features or endogenous processes (Kastner et al., 1998; Birn et al., 2006; Silver et al., 2007). Our predictions in higher-tier areas remain essentially unchanged if the source of symmetric biases on signal-strength were endogenous.

### Comparing brain measurements and computational models

RSA has been used to compare brain representations and network models, and taken to support Convolutional Neural Networks (CNNs) as models of human vision (Khaligh-Razavi and Kriegeskorte, 2014; Cichy et al., 2016). Recent work, however, highlights fundamental differences between how CNNs and humans represent visual information (Serre, 2019; Xu and Vaziri-Pashkam, 2021). We showed that a model that incorporates biological constraints, but does not compute anything, accounts for inconsistencies across studies due to different image statistics and RSA analysis choices. Incorporating similar constraints in CNNs, and considering the impact of the measurement process on RSA (Ramírez and Merriam, 2020), may help clarify these discrepancies.

### MVPA: Interpretational challenges and opportunities

MVPA encompasses numerous methods that consider the multivariate structure of neuroimaging signals when studying brain function (Haynes and Rees, 2006; Norman et al., 2006; Pereira et al., 2009; Tong and Pratte, 2012). Specifically, RSA aims to understand how information is represented in the brain by leveraging dissimilarities across experimental conditions (Kriegeskorte et al., 2008). Key challenges include how to estimate and normalize brain patterns, choose dissimilarity measure, and compare empirical and model DSMs. A growing literature aims to inform these choices and avoid interpretational pitfalls (Misaki et al., 2010; Diedrichsen et al., 2011; Smith et al., 2011; Garrido et al., 2013; Davis et al., 2014; Nili et al., 2014; Ramírez et al., 2014; Diedrichsen and Kriegeskorte, 2017; Ramírez, 2017; Cai et al., 2019; Ritchie et al., 2021; Kaniuth and Hebart, 2022). Here, we focused on the impact of mean subtraction across conditions and differences among angular and Euclidean pattern dissimilarity measures. While this study is not prescriptive, we have argued that angular and Euclidean distances provide fundamentally different descriptions of the data, and that angular distances are uniquely suited to reveal information regarding the form-of-tuning of indirectly measured neural populations (Ramírez, 2018). The simulations presented here help characterize differences and similarities in the behavior of three RSA variants as a function of low-level biases, given basic properties of the visual system, in the absence of a true underlying form of view-tuning.

## Conclusion

We proposed a parsimonious model considering feedforward projections, cortical magnification, interhemispheric crossings, and a measurement gain field. This model accounted for a range of unexplained similarities and inconsistencies found across fMRI-MVPA studies investigating facial viewpoint representations in humans. Our results suggest an important source of current disagreement may be due to analysis choices. We demonstrate how choice of distance measure and data-demeaning can lead to the impression that results across studies are inconsistent, when instead they offer different descriptions of the same process. Our results also suggest low-level feature imbalances across conditions may have been mistaken as evidence of mirror-symmetrically tuned neuronal populations. While this study is not prescriptive, it demonstrates the need for principled choices regarding data preprocessing, modeling, normalization, and pattern dissimilarity measure. Our results invite further methodological work to inform these choices and thus enable increasingly informative comparisons between empirical data and representational models. One way forward, we propose, resides in sufficiently constrained models able to robustly isolate meaningful pattern-components, possibly informative regarding the form-of-tuning of the underlying neuronal populations.

## ACKNOWLEDGEMENTS

We thank Charles Zheng and Daniel Handwerker for helpful feedback and suggestions. This work utilized the computational resources of the NIH HPC Biowulf cluster (http://hpc.nih.gov). This work was supported by the Intramural Research Program at NIMH (ZIAMH002783 and ZIAMH002966).

## Notes

### Competing Interest Statement

The authors have declared no competing interest.

