## Supplementary Material for "A unifying model for discordant and concordant results in human neuroimaging studies of facial viewpoint selectivity"

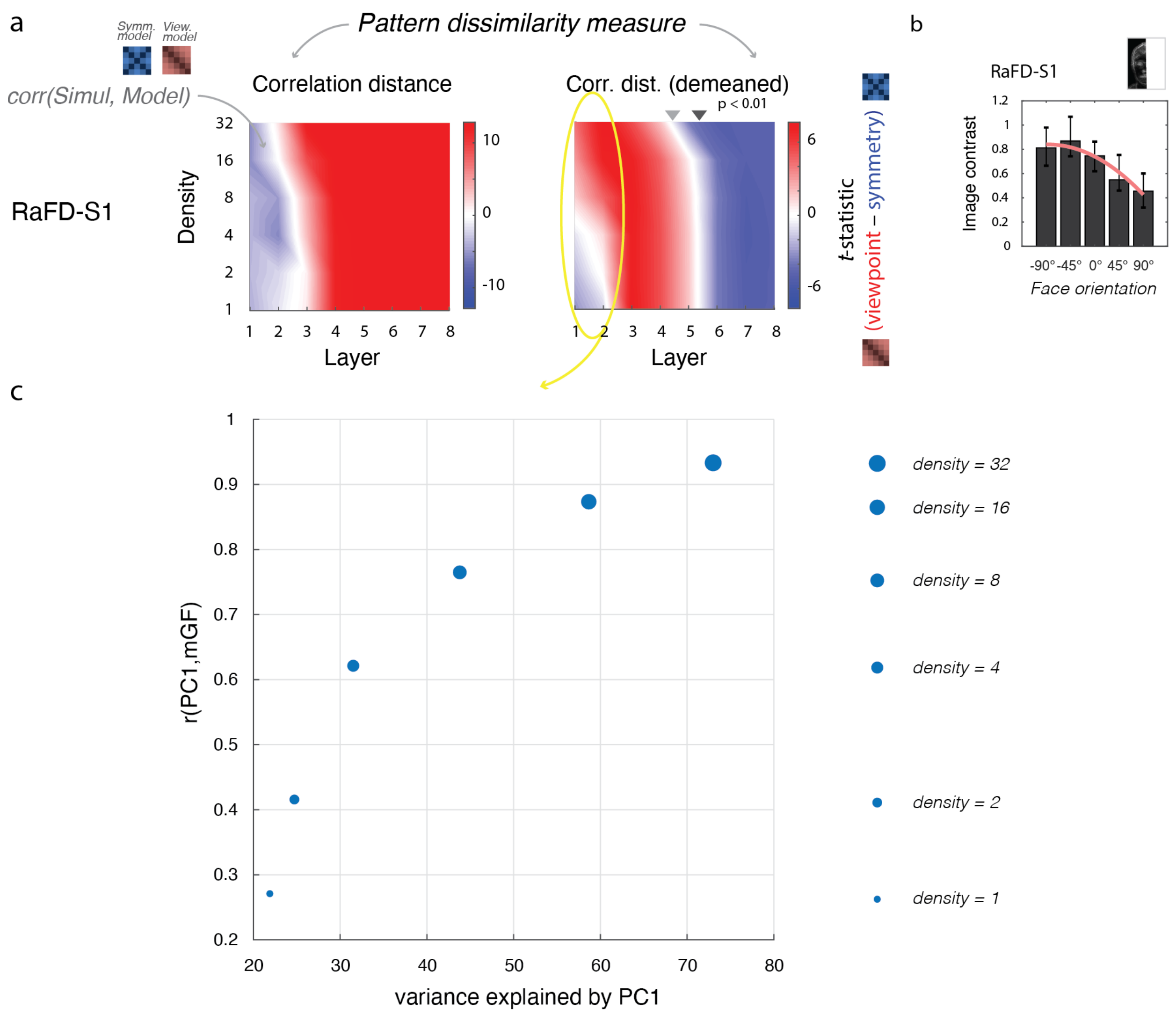


**Figure S1.** *Explanation of interaction between network density and form of view-tuning observed in network Layer 1 with $RSA_{corrDem}$ for the RaFD database and model-variant S1.*

*a*) The interaction between network density and form of view-tuning initially observed in Figure 8 is reproduced here and highlighted within a yellow ellipse. A bias to observe view-tuning with $RSA_{corrDem}$ is evident for early network layers and higher network densities (shown in red). In contrast, the opposite bias is observed for low densities, which reveal mirror-symmetry instead (shown in blue). As noted in the Discussion (see main text), this is due to an expected stronger influence for higher than lower network densities of the antisymmetric biases in signal strength observed across face views—the latter shown in panel *b*. If network density is high, after demeaning the simulated brain patterns across conditions, antisymmetry is induced on the anglular distances of the associated pattern-vectors due to the dominant antisymmetric trend observed in the pattern norms. This leads with $RSA_{corrDem}$ to a high association between the simulated data and the Viewpoint model. However, as can be seen in the leftmost plot showing $RSA_{corr}$ results—i.e., when the data are not demeaned—it turns out that the spatial structure of the images of this database also exhibits a genuine mirror-symmetric bias (revealed by the blue color observed in early network layers regardless of network density). If network density is low, this mirror-symmetric pattern component determines the nature of the impact of data demeaning, which in this case does *not* lead to observations of view-tuning with $RSA_{corrDem}$, but to the observation of mirror-symmetry. In other words, the impact of data demeaning depends on characteristics and the relative strength of the dominant pattern components, which we will next probe with Principal Component Analysis (PCA) to further clarify this matter.

c) *Relationship between network-density, variance explained by the 1$^{st}$ PC of patterns associated with the RaFD database, and the direction of the measurement Gain Field (mGF) in Layer 1.* For each network density-level (indicated by the width of the circular markers), the percentage of variance explained by the first PC is shown on the x-axis. PCA was computed on all patterns associated with all face orientations and identities for the RaFD database (S1 model-variant). The linear (Pearson) correlation between the 1$^{st}$ PC and the structure of the mGF is shown on the y-axis. As is evident in the graph, the higher the density, the more variance the 1$^{st}$ PC explains, and the more its spatial structure correlates with that of the mGF. Please note that the direction in multivariate voxel space associated with the mGF would explain virtually 100% of the variance for sufficiently high network densities. In this case, the energy of each face image (weighted by the cortical magnification function, and the hemifield representation) would determine its associated response pattern in network Layer 1. Information carried by the measured signal about the finer-scale spatial structure of the images would be negligible. As can be seen in the plot, if network-density is high ($d$=32), the 1$^{st}$ PC of the patterns associated with the images of this database is highly correlated with the direction defined by the mGF ($r$=0.95) and explains a large proportion of the variance (75%). The direction of the mGF is simply that of it's vector representation in voxel space. In this case, the antisymmetric imbalances in signal-strength across face-views noted in half-images and early network layers (see panel b) will determine the influence of data demeaning on pattern-vector angles. By the contrary, if network density is low ($d$=1), then mirror-symmetric biases identified in the spatial structure of the images will instead dictate the nature of the influence of mean subtraction. As expected, in this latter case the correlation of the 1$^{st}$ PC with the mGF is low ($r$=0.28) and only explains 22% of the variance. The mirror symmetric biases revealed with $RSA_{corr}$ in panel *a*, leftmost plot, will in this case mostly determine the influence of data demeaning on pattern-vector angles. In contrast to higher network densities, lower densities lead with $RSA_{corrDem}$ to observations biased towards finding mirror-symmetry. This phenomenon is closely related to the influence of measurement scale on $RSA_{Euc}$ and $RSA_{corr}$ described in Ramírez (2018, Figure 6). For more on the impact of the measurement process, data granularity, and mean subtraction on RSA, please see: (Ramírez et al., 2014, 2020; Ramírez, 2017; Ramírez and Merriam, 2020).

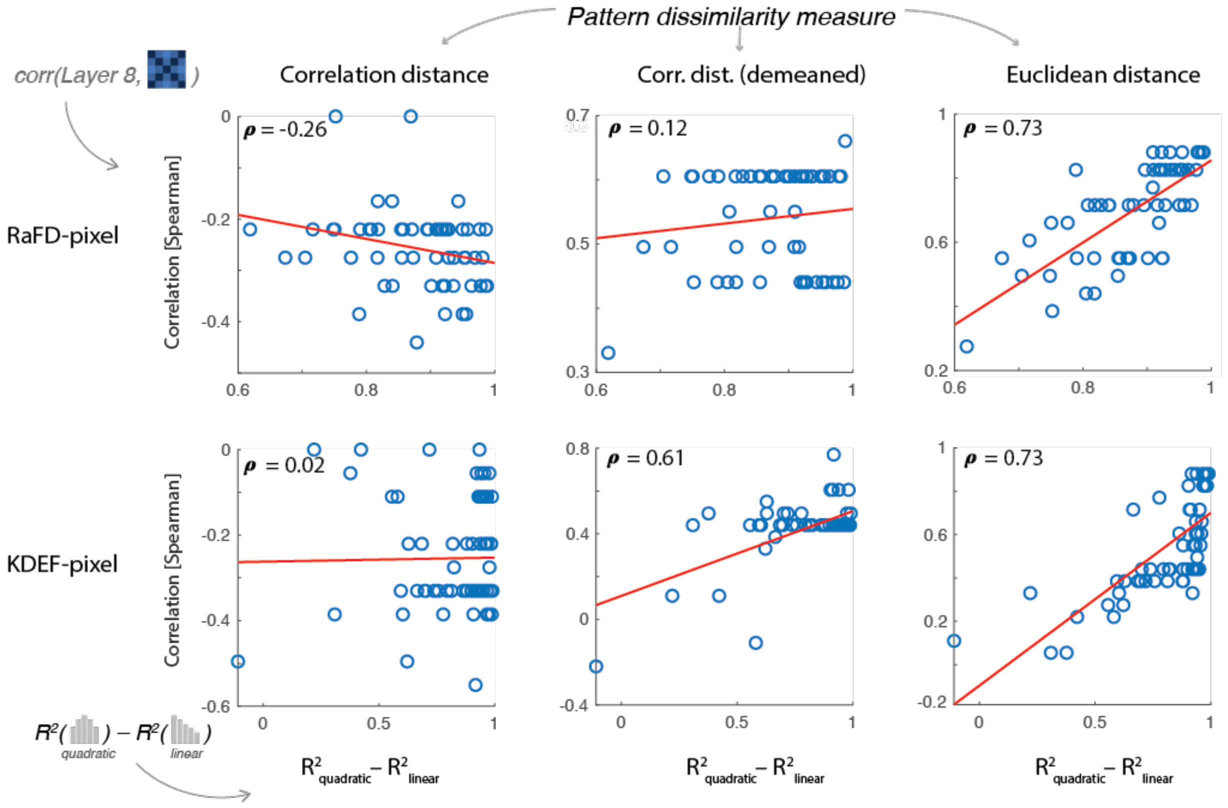


**Figure S2.** *Accounting for mirror-symmetry in RSA for single face-identities: pixel-level analysis.* Layout and analyses as in Figure 9. Here, however, results from the pixel-level model are shown instead of S1-level model. As observed for S1-level results, the difference in $R^2$ between the quadratic and linear components of the layer 8 norm-profile of each face identity (x-axis) is significantly positively correlated with the degree of mirror-symmetry (y-axis) observed with $RSA_{Euc}$ (both RaFD and KDEF: $r = 0.73$, $p < 0.001$). $RSA_{corrDem}$ results exhibit qualitatively similar trends (RaFD: $r = 0.12$, *n.s.*, KDEF: $r = 0.61$, $p < 0.001$). Note that while the sign of the observed association coincides with that observed with model-variant S1 (cf. Figure 9), the correlation observed here for RaFD is weak and not statistically significant. Critically, however, and consistent with the logic of our argument, the association between the difference of $R^2_{quad}$ and $R^2_{lin}$ and $RSA_{Corr}$ with the mirror-symmetric model is negative or close to zero, and in both cases statistically insignificant (RaFD: $r = -0.26$, KDEF: $r = 0.02$, both $p > 0.05$).

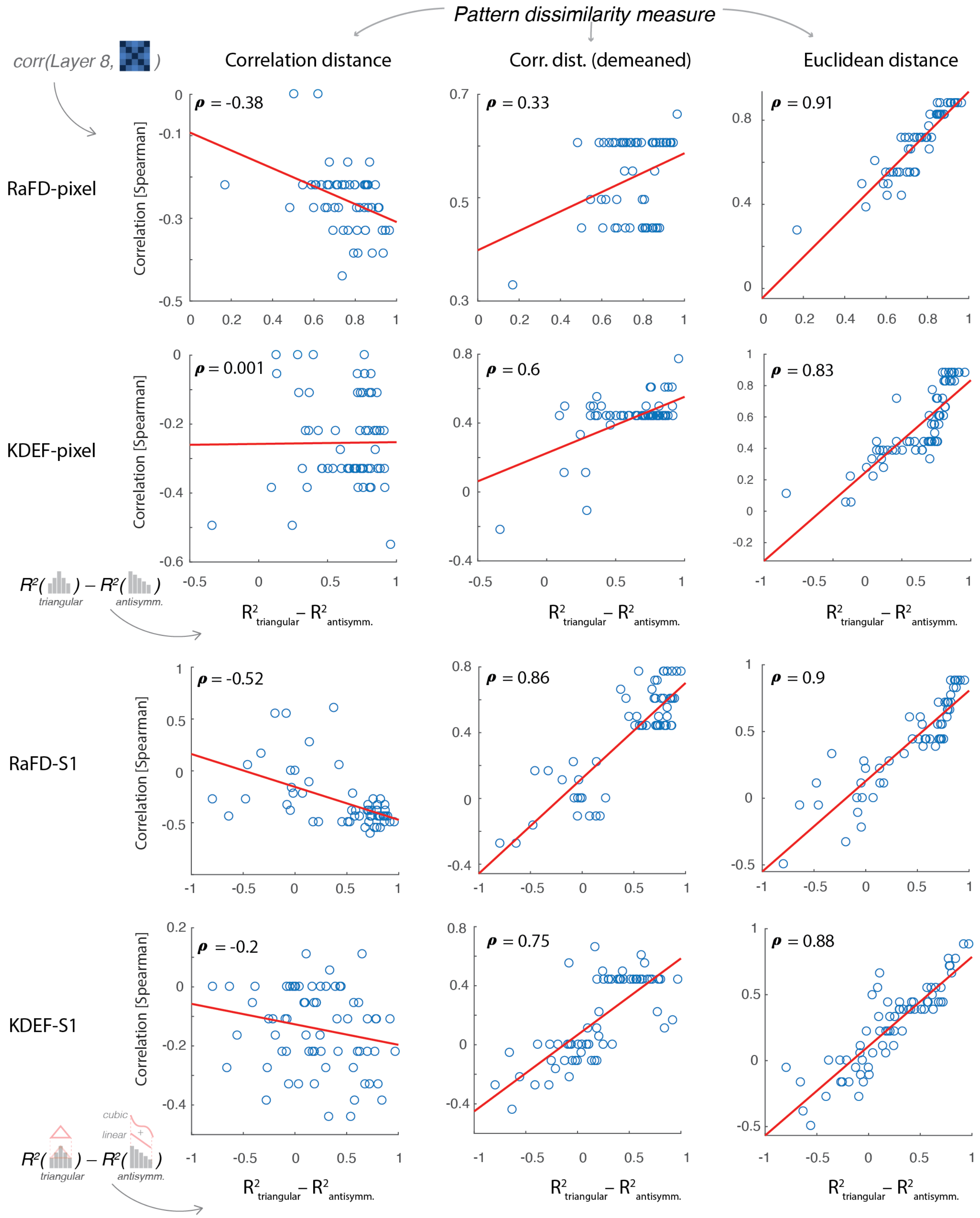


**Figure S3.** *Further accounting for mirror-symmetry in RSA for single face-identities.* Layout and analysis as in Figure 9. Here, however, results for both the pixel- and S1-level model variants are shown, and a different decomposition of the norm-profiles explored. Instead of using constant, linear, and quadratic regressors, as in Figures 9 and S2, we relied on a decomposition consisting of constant, symmetric (now modeled with a triangular regressor, see inset in Figure S3, bottom left corner) and antisymmetric components. The antisymmetric component consisted of variance explained by linear and cubic polynomial trends. We reasoned that including a triangular (symmetric) regressor and a more flexible antisymmetric regressor might better predict the degree of mirror symmetry for the Euclidean distance ($RSA_{Euc}$) and possibly also $RSA_{corrDem}$. This expectation was based on two reasons: (i) a triangular shape changes linearly between each profile view and the frontal view, unlike the quadratic trend, and (ii) a more general antisymmetric regressor captures variance unaccounted by the linear regressor. As observed for S1-level results, the difference in $R^2$ between the quadratic and linear components of the layer 8 norm-profile of each face identity (x-axis) significantly positively correlates with the degree of mirror-symmetry (y-axis) observed with $RSA_{Euc}$. As expected, this association is stronger than observed with the previous decomposition in 4/4 analyses (RaFD and KDEF: $r$ = [0.83, 0.91], all $p < 0.001$). $RSA_{CorrDem}$ trends were consistent in direction with the original decomposition in Figures 9 and S2 and always statistically significant (RaFD-pixel: $r = 0.33$, p = 0.013; KDEF-pixel: $r = 0.60$, $p < 0.001$; RaFD-S1: $r = 0.86$, p < 0.001; KDEF-S1: $r = 0.75$, $p < 0.001$). The observed associations outperformed the previous decomposition, however, in only 2/4 analyses. Most importantly, and consistent with the logic of our argument, the association between the difference in $R^2$ of the new symmetric and antisymmetric components and $RSA_{Corr}$ with the mirror-symmetric mDSM were either significantly *negative*, or statistically insignificant (RaFD-pixel: $r = -0.38$, p = 0.003; KDEF-pixel: $r = 0.01$, $p = 0.94$; RaFD-S1: $r = -0.58$, p < 0.001; KDEF-S1: $r = 0.01$, $p = 0.94$).

### Supplementary Discussion

#### *A. On the weak mirror-symmetric effect observed in a study that jointly analyzed left and right FFA voxels*

The study by Guntupalli et al. (2017) conducted RSA on uncentered data. These authors explored two different distance measures. For searchlight analyses, they used classification accuracies from pairwise linear SVM classifiers—sensitive to the Euclidean distance between the endpoint of pattern vectors. These searchlight analyses revealed evidence of view-tuning throughout occipitotemporal cortex, bilaterally, and no evidence of mirror-symmetry (see Discussion, in main text). When jointly analyzing voxels from both cerebral hemispheres with $RSA_{corr}$, however, Guntupalli et al. observed not only evidence of a view-tuned representation, as in the first analysis, but now also evidence in FFA of a weaker mirror-symmetric effect. These observations may seem partially inconsistent with our model. Three factors, however, may help explain these findings. First, Guntupalli et al. used a one-back repetition detection task that did not require subjects to fixate on a known image location. If we assume these researchers observed in EVC the same anti-symmetric pattern of responses across views found by Axelrod and Yovel (2012) (Experiment 1) when relying on the same task, then a simple explanation emerges. A strong anti-symmetric signal across views in EVC would explain the generalized lack of mirror-symmetry noted with their searchlight analysis (~$RSA_{Euc}$). Indeed, Axelrod and Yovel reported markedly reduced symmetric trends on classification rates in FFA in their Experiment 1 (one-back task, free viewing) compared with Experiment 2 (color-change detection of fixation spot). A second factor is that for FFA analyses with $RSA_{corr}$, unlike searchlight analyses, Guntupalli et al. pooled voxels from both cerebral hemispheres. A plausible source of the weak mirror-symmetric effect observed for the $RSA_{corr}$ analysis are previously reported differential responses across hemispheres that interact with face-view (Hung et al., 2010; Caharel et al., 2011; Rossion and Caharel, 2011; Verosky and Turk-Browne, 2012; Or et al., 2021). Providing evidence of mirror-symmetry separately in each cerebral hemisphere would help rule-out this possibility. Overall, our analyses on uncentered data match observations in experiments requiring subjects to fixate on a central image location. The pattern of average activations for faces in different viewpoints in EVC reported by Axelrod and Yovel (2012) for the one-back repetition task (due to systematic eye movements towards specific facial features) provides a straightforward account of RSA observations in the two experiments that relied on this task. The third factor relates to the possible influence of signal-to-noise effects on RSA. Such effects have been shown to possibly lead to observations of weak "mirror symmetry" in the complete absence of mirror-symmetrically tuned neurons, as detailed in Ramírez (2018).

*B. More on CNNs and the importance of incorporating biologically-inspired constraints*

Previous neuroimaging work showed the importance of cortical magnification to account for univariate response profiles to faces shown in various views in EVC and FFA (Yue et al., 2011). Our observations further suggest that CM and interhemispheric crossings are important constraints for networks aiming to model multivariate measurements of the primate visual system. Despite their success in solving complex visual tasks, Convolutional Neural Networks (CNNs) (Güçlü and Gerven, 2015; Kriegeskorte, 2015; Cichy et al., 2016; Kubilius et al., 2016; Xu and Vaziri-Pashkam, 2021) do not usually consider these constraints (but see Larochelle and Hinton, 2010; Costa et al., 2021; Wang et al., 2021). Doing this may help CNNs achieve more human-like performance and render them more easily comparable to brain measurements. Interestingly, Cichy et al. (2016) explored CNNs trained in a categorization task, and compared them with networks trained with either randomly labeled, or smoothed noise images. All three networks exhibited hierarchical structure that correlated with human neuroimaging data, but the CNN trained on categorization best explained the data. These results suggest hierarchically organized randomly-connected networks like the one we constructed and explored in this study, as well as CNNs trained with randomly-labeled images, as well as smoothed noise images, can reveal meaningful properties that systematically relate to neuroimaging data.
